# Titrating CD47 by mismatch CRISPRi reveals incomplete repression can eliminate IgG-opsonized tumors but CD47 heterogeneity limits induction of anti-tumor IgG

**DOI:** 10.1101/2022.09.27.509740

**Authors:** Brandon H. Hayes, Hui Zhu, Jason C. Andrechak, Dennis E. Discher

**Affiliations:** Molecular and Cell Biophysics Lab, University of Pennsylvania, Philadelphia, PA, USA; Physical Sciences-Oncology Center at Penn, University of Pennsylvania, Philadelphia, PA, USA; Bioengineering Graduate Group, University of Pennsylvania, Philadelphia, PA, USA

## Abstract

Phagocytic elimination of solid tumors is an attractive mechanism for immunotherapy – particularly because of the possible induction of anti-cancer immunity. The phagocytic potential of macrophages is limited, however, by the CD47-SIRPα checkpoint, and how much CD47 disruption is needed for efficacy remains unclear, even when tumors are opsonized by a pro-phagocytic antibody. Here, CRISPR-interference (CRISPRi) is applied with a large set of sgRNAs to produce a broad range of CD47 knockdowns in B16F10 melanoma, which is generally found to be resistant to the heavily studied PD-1 blockade. Guided by 3D immuno-tumoroid results, we identify a critical CD47 density below which macrophage-mediated phagocytosis dominates proliferation in the presence of an otherwise ineffective pro-phagocytic antibody (anti-Tyrp1). Growing tumors and immuno-tumoroids generally show selection for CD47-positive cells, but some mice reject tumors having >97% mean repression of CD47 or even having 80% repression – unless mixed with 50% repressed cells. Interestingly, long-term survivors have *de novo* pro-phagocytic IgG antibodies that increase in titer with depth of repression and also with early accumulation of tumor macrophages. Given well-known limitations of antibody permeation into solid tumors, our studies set a benchmark for anti-CD47 blockade and suggest deep disruption favors acquired immunity.

## Introduction

Macrophages are increasingly recognized as attractive effector cells for cancer immunotherapies due to their ability to recognize and destroy cells through phagocytosis (Alvey & Discher, 2017; Klichinsky et al., 2020; Lee et al., 2016; Morrissey et al., 2018). The efficacy of phagocytosis is generally inhibited by the CD47-SIRPα axis, in which CD47 expressed on “self” cells (including cancer cells) interacts with SIRPα on macrophages (Okazawa et al., 2005; Oldenborg et al., 2000). Generally, tumor cell engulfment by macrophage-mediated phagocytosis can occur when using tumor antigen-targeting monoclonal antibodies that engage Fc receptors on macrophages (Alvey et al., 2017; Suter et al., 2021). Such therapies can improve efficacy when paired with anti-CD47 antibody blockade in lymphoma patients (Advani et al., 2018), whereas disabling *only* the CD47-SIRPα axis between macrophages and cancer cells does not appear effective in the clinic (Jalil et al., 2020) and remains controversial even in syngeneic mouse model experiments (Horrigan 2017) designed to reproduce initial reports (Willingham et al., 2012). Even if combinations with tumor antigen-targeting antibodies or other pro-phagocytic cues prove essential to efficacy, the levels of CD47-SIRPα disruption that are needed have remained unclear.

While we have recently reported that CRISPR/Cas9-mediated CD47 knockout can lead to tumor rejection and durable cures when paired with opsonizing IgG (Dooling et al., 2022), complete knockout does not currently represent clinically viable strategies for patient tumors. *In vivo* genome editing also may be not effective on all desired target cells (Mout et al., 2017), which creates significant heterogeneity that may ultimately lead to recurrence. Furthermore, there are ongoing concerns about both the delivery feasibility, efficacy and safety of CRISPR/Cas9 knockout in patients (Lino et al., 2018; Wilbie et al., 2019; You et al., 2019). For example, double strand breaks during the genome editing can result in genotoxicity, induce chromothripsis, lead to the expression of oncogenic fusion proteins, or give rise to dysregulated gene expression (Azangou-Khyavy et al., 2020; Cullot et al., 2019; Kosicki et al., 2018; Leibowitz et al., 2021). Lastly, knockout studies do not realistically represent clinical situations in which complete inhibition by anti-CD47 blockade is not possible due to low permeation of solid tumors (Ingram et al., 2017).

CRISPR-interference (CRISPRi), however, can address several of these concerns. This system uses a catalytically inactive dCas9 that does not induce double strand breaks, minimizing off-target or genotoxic effects (Gilbert et al., 2014). CRISPRi also enables simulation of more realistic “intermediate” inhibitory levels of CD47 achievable in bulk cancer cell populations by using base pair mismatches to titrate sgRNA efficacy. This also enables us to explore unanswered questions concerning to what extent CD47 expression heterogeneity can lead to cancer cell escape from immune response. Myeloid leukemias have previously been found to adopt the physiological response of upregulating CD47 that hematopoietic stem and progenitor cells use during an insult that causes mobilization (Jaiswal et al., 2009). CD47 heterogeneity expression may induce macrophage-mediated pressure for CD47-high expressors (cancer and progenitors alike) that can eventually gain survival advantage and induce recurrence.

To this end, we hypothesized that mismatch-CRISPRi (Jost et al., 2020; Hawkins et al., 2020) would reveal a critical CD47 density at which therapeutic clearance of tumors by macrophages could be obtained without requiring complete ablation of CD47. Our results identify levels of repression that achieve therapeutic clearance, but they also highlight that both CD47 expression heterogeneity and incomplete repression affect the titer of *de novo* anti-cancer IgG and possibility of cancer recurrence.

## Results

### Mismatched sgRNAS in CRISPRi titrate CD47 expression with broad heterogeneity

To identify critical CD47 densities at which phagocytosis starts showing inhibited activity, we used a recently described CRISPRi method (Jost et al., 2020) that uses sgRNAs with base pair mismatches to genomic DNA to affect binding of a nuclease dead Cas9 variant (dCas9) (**Fig. 1A**). We used our mismatch-CRISPRi in B16F10 mouse melanoma model because it responds poorly to T cell checkpoint blockade and CD47-SIRPα blockade/disruption *in vivo* (Sockolosky et al., 2016). Furthermore, the B16F10 model expresses Tyrp1 antigen, which can be targeted by a therapeutic IgG antibody that promotes phagocytic uptake by macrophages (Andrechak et al., 2022; Hayes et al., 2020; Khalil et al., 2016).

**Figure 1.**
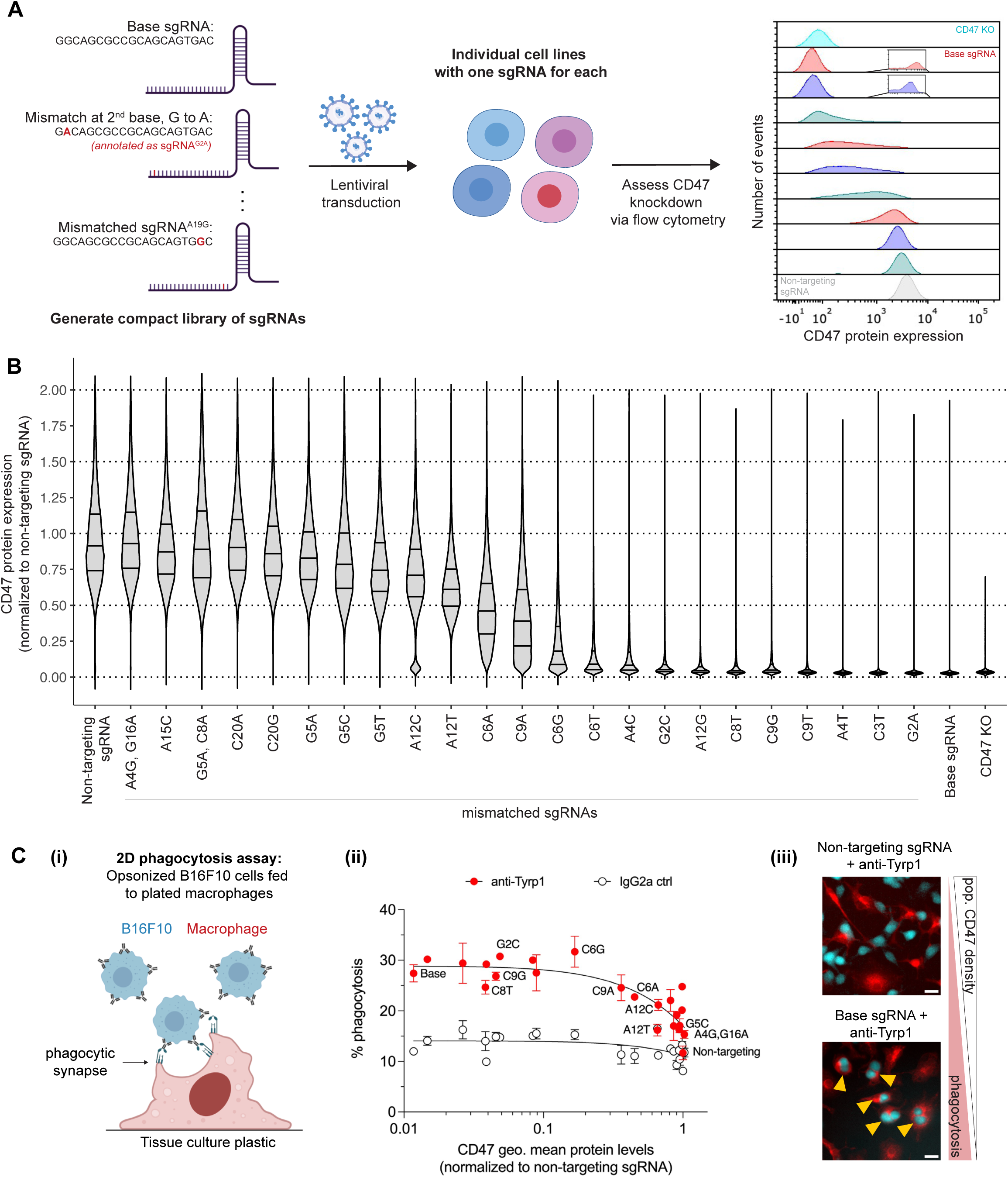
CRISPRi with mismatched sgRNAs effectively titrates CD47 expression in cancer cells and generates populations with broad CD47 heterogeneity. **(A)** Schematic illustrating the development of a compact library of mismatched sgRNAs used with the CRISPR interference (CRISPRi) system to induce knockdown. An initial base sgRNA (from Jost et al., 2020) was modified by performing either one or two base pair mismatches at a particular nucleotide position of interest. For nomenclature, each sgRNA is annotated with its original base at the nucleotide position of interest (from the 5’-end), followed immediately by the new base for the mismatch. Example: Switching the second base in the parental sgRNA, originally a guanine (G), to adenine (A) leads to sgRNA^G2A^. sgRNAs were then packaged into lentivirus and delivered to target cells. A single sgRNA was delivered per titrated cell line. After antibiotic selection to kill non-transduced cells, all developed CD47-titrated cell lines had their CD47 protein expression levels measured by flow cytometry. **(B)** Distribution of CD47 levels in B16F10 mouse melanoma cells using a perfectly matched (base) sgRNA, a subset of mismatched sgRNAs, and a non-targeting sgRNA. Additionally, a B16F10 cell line (from the same parental line) with regular CRISPR/Cas9-mediated knockout of CD47 is included for comparisons to complete ablation. All data are obtained by flow cytometry and normalized to the geometric mean CD47 expression of the B16F10 population transduced with the non-targeting sgRNA. **(C) (i)** Schematic of phagocytosis of CD47-titrated B16F10 cell lines by bone marrow-derived macrophages, with or without anti-Tyrp1 for opsonization on two-dimensional tissue culture plastic. **(ii)** Quantification of macrophage-mediated phagocytosis on 2D plastic. B16F10 cells with increasingly deeper knockdown of CD47 are more readily phagocytosed by bone-marrow derived macrophages when therapeutically opsonized with IgG. Phagocytic activity, however, plateaus starting at ∼20% of the original density and remains at similar levels even with deeper knockdown of CD47. **(iii)** Images on far-right are representative of phagocytic activity against B16F10 cells with wild-type levels of CD47 (non-targeting sgRNA) and deep knockdown (∼99% depletion). Scale bar = 20 μm.

First, we identified a base sgRNA (Horlbeck et al., 2016) that deeply represses CD47 (by 97-99% of original levels) (**Fig. S1A, S1B**). While this base sgRNA induces deep repression, we do note that there is still a small subset of cells (∼1% of the entire population) that still maintain intermediate-to-high CD47 levels (**Fig. 1A – flow cytometry plot insets; Fig. S1A**). Nonetheless, the deep repression enables us to systematically perform nucleotide mismatches to greatly alter sgRNA efficacy to generate the desired titration. For choosing mismatches, we assumed that nucleotide position mattered more than the nucleotide itself, based on previous computational modeling experiments (Boyle et al., 2017; Eslami-Mossallam et al., 2022; Jones et al., 2021). We then generated an initial compact sgRNA library with mismatches at nearly all nucleotide positions to identify possibly sensitive base sites relative to the protospacer adjacent motif (PAM) (**Fig. S1C**). Like previously reported results (Jost et al., 2022), we generally found that mismatches near the PAM strongly attenuated sgRNA efficacy. Conversely, mismatches in the PAM-distal region showed little effect. Mismatches in the middle region produced the largest variability of knockdown, in terms of repression efficacy and CD47 expression heterogeneity. We thus identified a subset of sgRNAs with mismatches at different nucleotide positions that covered deep, intermediate, and little repression of CD47 and performed the remaining nucleotide mismatches (**Fig. S1D**). In addition, we noticed that cell-to-cell variation in expression for a given sgRNA tends to be highest in the intermediate range of expression, particularly in the sigmoidal transition range, which is consistent with the fundamentals of binding and binding capacity (**Fig. S2**). The final sgRNAs used for subsequent experiments and their CD47 knockdown distribution are shown in **Fig. 1B**.

### CD47 titration identifies critical density limits that modulate phagocytosis

We hypothesized that macrophage-mediated phagocytosis of B16F10 would be maximized with the base sgRNA that induces deep CD47 repression and that decreasing levels of knockdown efficacy would eventually hinder macrophage phagocytic potential as more CD47 is expressed. To identify if there are any target cell critical CD47 densities at which this phagocytic activity shows sensitivity to CD47-SIRPα checkpoint, we first performed conventional 2D phagocytosis assays in which cancer cell suspensions are added to immobilized bone marrow-derived macrophages (BMDMs) (**Fig. 1C**). Prior to feeding the cancer cell suspension to BMDMs, the B16F10 cells were treated with either anti-Tyrp1 for IgG opsonization to stimulate phagocytosis or mouse IgG2a isotype control. The base sgRNA leads to a ∼2.5-fold increase in phagocytic uptake when B16F10 are opsonized with anti-Tyrp1 compared to the mouse IgG2a control. We confirmed that CRISPRi repression did not increase Tyrp1 levels. SIRPα levels increased (**Fig. S3**), consistent with previous studies on CD47-SIRPα *cis* interactions (Hayes et al., 2020) showing that reduced CD47 surface density frees up more SIRPα molecules on that same cell surface. These results are also consistent with previous studies on the importance of opsonizing antibodies that engage Fcγ receptors (Bakalar et al., 2018; Morrissey et al., 2020) to ultimately induce a phosphorylation cascade that activates actomyosin tension to facilitate phagocytic internalization (Tsai & Discher, 2008).

Our mismatch-CRISPRi titration of CD47 further showed gradual reductions in macrophage-mediated phagocytosis of cancer cells as sgRNA efficacy decreased. All sgRNAs that repressed CD47 by 80% or more showed nearly identical levels of cancer cell phagocytosis. Once sgRNA knockdown efficacy was lower than ∼80%, we noted a downward shift in phagocytic activity (**Fig. 1C**). These results support the overall hypothesis that macrophages are sensitive to the overall surface density of CD47 molecules on target cancer cells and that tuning these levels yields a distribution of phagocytic activity.

To model the biophysical properties of the 3D tumor microenvironment and proliferative capacity of tumor masses, we then engineered “tumoroids” of the B16F10 cells in non-adhesive culture plates (**Fig. 2A-B**). We chose cultures with non-targeting sgRNA (wild-type CD47 levels), base sgRNA (deep repression), and 10 other sub-lines that covered a broad range of knockdown efficacy and overall heterogeneity for these tumoroid studies. BMDMs were added to pre-assembled tumoroids at a 3:1 ratio with either anti-Tyrp1 or mouse IgG2a control (**Fig. 2C-D**). We observed that B16F10 tumoroids with anywhere from 64-97% CD47 repression were all successfully cleared by macrophages when opsonized with anti-Tyrp1 (**Fig. 2C, 2D; S4A**). The sgRNA^C6G^ variant fits within this grouping whereas the sgRNA^C6A^ variant did not. This distinction became clear only at 48 h of the co-cultured tumoroids but was unclear at 24 h and also unclear in 2D phagocytosis assays (**Fig.1C-i**). We further found that knockdown levels of roughly 50% managed to suppress tumoroid growth when opsonized but could not fully clear all tumoroids (**Fig. 2C-D; Fig. S4B**). Knockdown of 35% and lower failed to clear any tumoroids or even inhibit their growth (**Fig. 2C-D; Fig. S4C**). Altogether, these immuno-tumoroid studies further highlight a CD47 surface density regime at which macrophage-mediated phagocytosis *starts* to become inhibited. A Pearson correlation analysis of the 2D assays and 3D tumoroid results also shows a strong correlation between both methodologies, suggesting 2D assays can predict 3D tumoroid success (**Fig. S5**).

**Figure 2.**
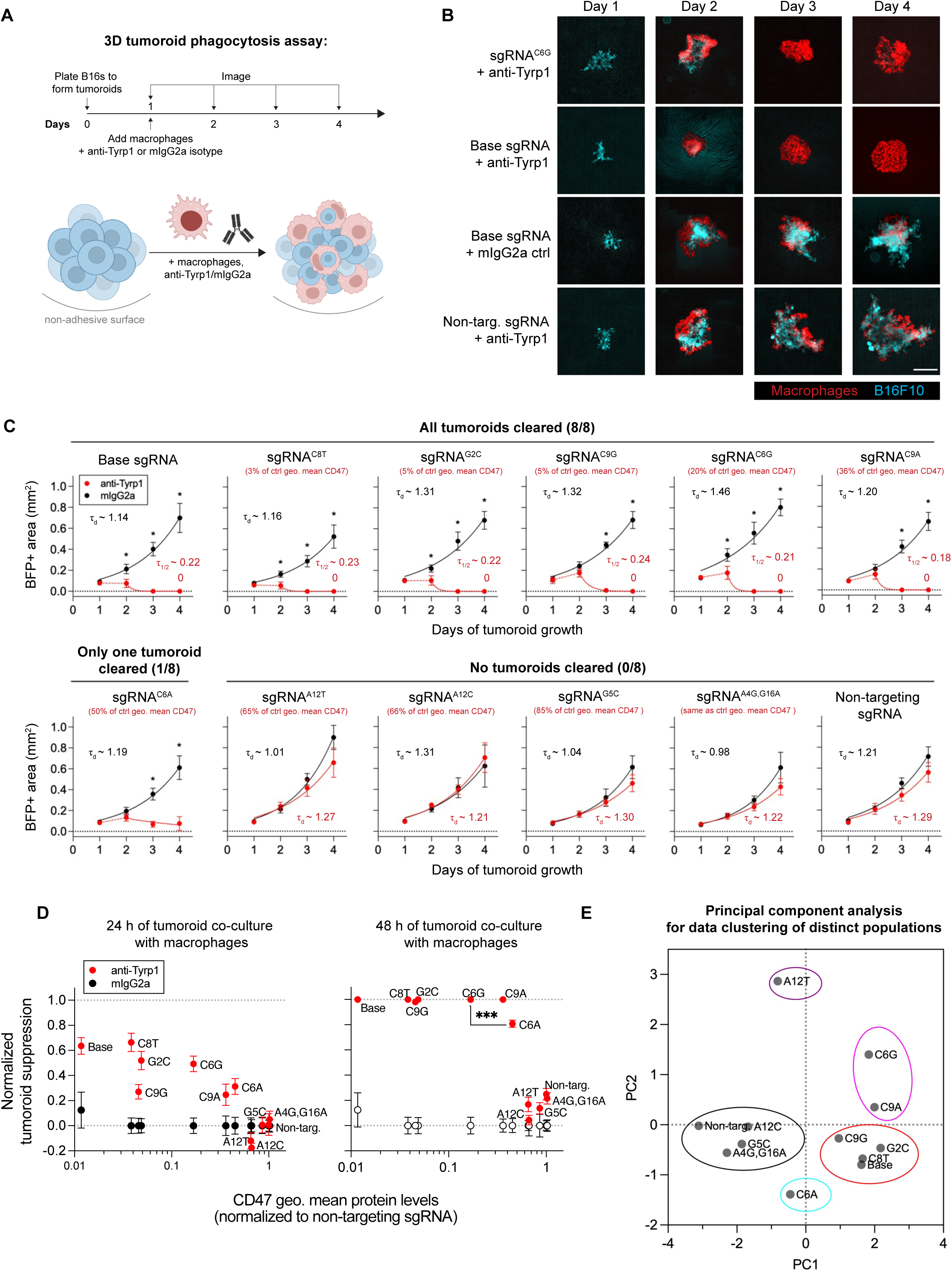
Incomplete but significant repression of CD47 drives clearance of tumoroids. **(A)** Timeline and schematic for generating engineered “immuno-tumoroids” for time-lapsed studies of macrophage-mediated phagocytosis of cancer cells. Tumoroids are formed by culturing B16F10 cells on non-adhesive surfaces in U-bottom shaped wells. ∼24 h after plating, bone marrow-derived macrophages with or without opsonizing anti-Tyrp1 are added to the cohesive B16F10 tumoroid. Immuno-tumoroids are imaged at the list timepoints. **(B)** Representative fluorescence images depicting either growth or repression of B16F10 cells (light blue) in immuno-tumoroids from days 1-4. Bone marrow-derived macrophages, shown in red, were added at a 3:1 ratio to initial B16F10 number after the day 1 images were acquired. Scale bar = 0.5 mm. **(C)** Tumoroid growth was measured by calculating the BFP+ area at the indicated timepoints (mean *±* SD, n = 8 total tumoroids from two independent experiments for each sgRNA). Opsonization of target B16F10 cells with anti-Tyrp1 limits growth of immuno-tumoroids only after at least ∼64% of the original CD47 density is depleted. After this critical knockdown threshold is reached, macrophages can clear tumoroids regardless of remaining CD47 levels. Together, 3D tumoroid and 2D phagocytosis assays suggest that repression to ∼20-36% of the original CD47 density levels is sufficient for therapeutic response with IgG opsonization. Statistical significance was calculated by the Mann-Whitney test (unpaired, two-tailed, * p < 0.05) comparing the BFP+ area between conditions with or without anti-Tyrp1 at the indicated timepoint. **(D)** Quantification of normalized tumoroid suppression 24 h (left) and 48 h (right) after addition of macrophages to tumoroids (mean *±* SD, n = 8 total tumoroids from two independent experiments for each sgRNA). Top dashed line indicates complete suppression, while bottom dashed line indicates no suppressive effect. A\Statistical significance between tumoroids made of sgRNA^C6G^ and sgRNA^C6A^ was calculated by an unpaired two-tailed t-test with Welch’s correction (* p < 0.05). **(E)** Principal component analysis (PCA) of the cell lines with the sgRNAs used in all phagocytosis and tumoroid experiments. The following characteristics of each cell line were used: geometric mean of CD47 expression, geometric standard deviation of CD47 expression, overall tumoroid clearance percentage, *in vitro* tumoroid growth rate with and without anti-Tyrp1, *in vitro* 2D phagocytosis percentages with and without anti-Tyrp1. PCA analysis identifies different data clusters from which to choose B16F10 cell lines for future *in vivo* experiments.

### Significant but incomplete repression of CD47 leads to tumor rejection and *de novo* IgG

While previous studies (Andrechak et al., 2022; Dooling et al., 2022; Li et al., 2020) show that complete CD47 ablation can favor rejection of B16F10 tumors, we further sought to find intermediate-CD47 expression levels that could yield similar rejection while being more clinically realistic. To select this intermediate CD47-expressing cell line for *in vivo* xenografts, we performed principal component analysis (PCA) to cluster cell lines with different knockdown levels (different sgRNAs) (**Fig. 2E**). We found that B16F10 cell lines with ∼20 and 36% knockdown (sgRNA^C6G^ and sgRNA^C9A^) uniquely clustered together, separately from the base sgRNA which we initially intended to use as a control. To maximize the possibility of tumor rejection, we chose with B16F10 with sgRNA^C6G^ since it had deeper repression while still showing decent variability in knockdown efficacy (**Fig. 1B**). We also generated a mosaic CD47 knockdown line by mixing B16F10 with sgRNA^C6G^ and sgRNA^C6A^ (**Fig. S6**). Although repression show a distribution of knockdown that includes CD47-intermediate and CD47-high cells, we hypothesize that an existing significant population of CD47-low cells is sufficient to stimulate anti-cancer macrophage activity that can aid in overall tumor suppression. Generating this mosaic culture allowed us to probe another intermediate CD47-expressing cell line while maintaining a significant CD47-low population to potentially activate immune response.

We proceeded to establish tumors in mice with B16F10 cells with wild-type CD47 levels (non-targeting sgRNA) (**Fig. 3A-left**), deep repression (base sgRNA) (**Fig. 3A-right**), and intermediate CD47-expression levels (sgRNA^C6G^ and mosaic) (**Fig. 3B**). For the wild-type and deep knockdown tumors, we separated mice into groups that received either anti-Tyrp1 or mouse IgG2a isotype control. Anti-Tyrp1 treatment seemingly eliminated tumors with deep CD47 knockdown in 23% of mice for at least 100 days, whereas control tumors (including mouse IgG2a-treated deep CD47 knockdown and both anti-Tyrp1-treated and mouse IgG2a-treated tumors) all exhibited exponential growth and no survivors (**Fig. 3A, 3C**). Given there were no survivors with mouse IgG2a regardless of knockdown level, like previous studies (Dooling et al., 2022), intermediate CD47-expressing tumors only received anti-Tyrp1. Tumors with sgRNA^C6G^ also had 30% mice surviving after 100 days when treated with anti-Tyrp1 (**Fig. 3B-bottom**), confirming that complete inhibition or ablation of CD47 is not required for tumor rejection. The mosaic tumors, however, all grew exponentially (**Fig. 3B-top**), suggesting that even with knockdown, CD47 expression heterogeneity is an important factor in tumor rejection and most likely requires a majority of the tumor to be CD47-low.

**Figure 3.**
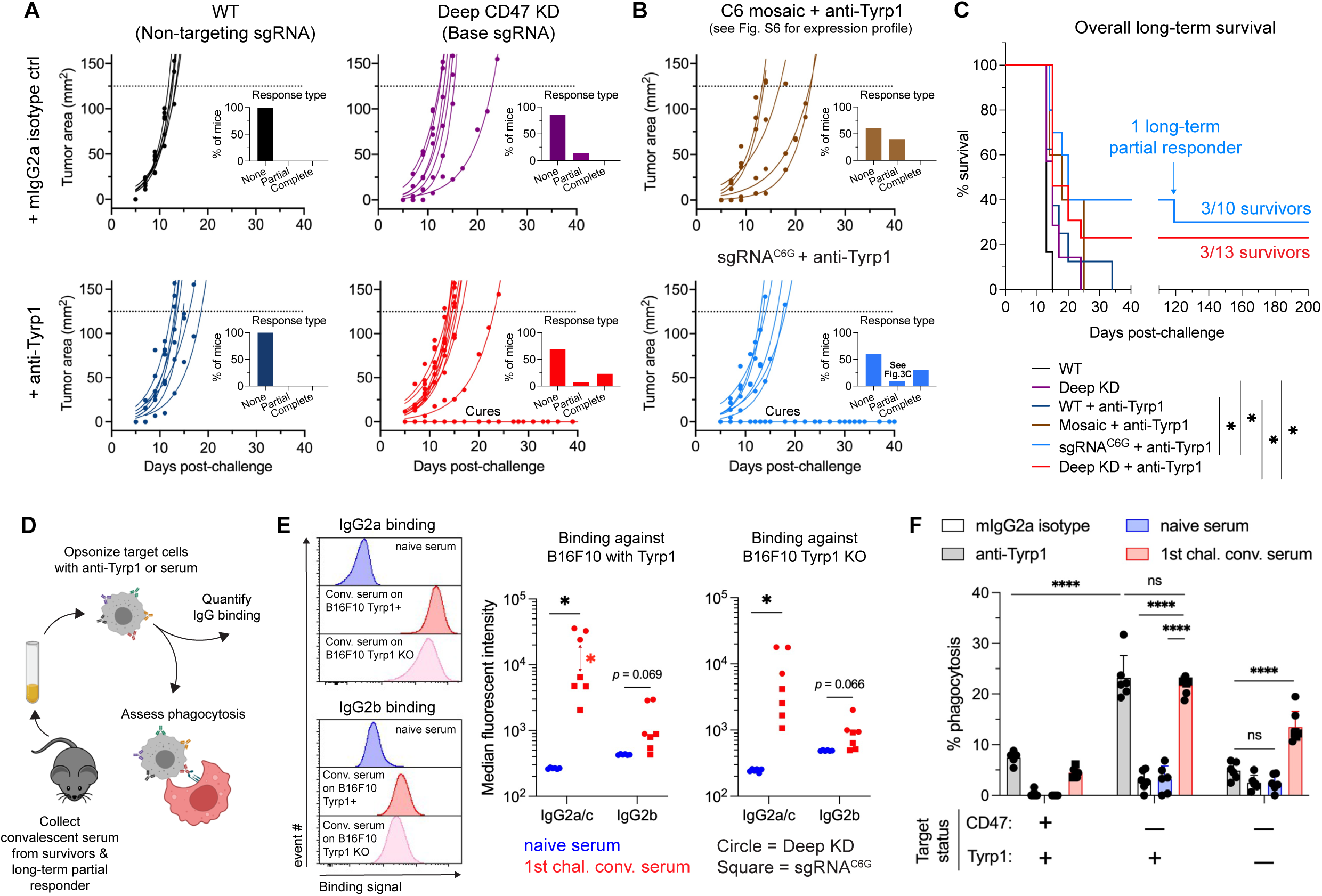
Suppression of CD47 expression to critical density or lower represses syngeneic tumors *in vivo* and leads to anti-tumor antibody generation. **(A)** Tumor growth curve of projected tumor area versus days after tumor challenge, with B16F10 cells expressing wild-type levels of CD47 (abbreviated at WT) or near complete knockdown (base sgRNA). Each line represents a separate tumor and is fit with an exponential growth equation: A = A_0_e^kt^. Complete anti-tumor responses in which a tumor never grew are depicted with the same symbol as their growing counterparts and with solid lines at A = 0. For experiments with B16F10 with the non-targeting sgRNA, 6 mice received mouse IgG2a isotype control and 8 mice received anti-Tyrp1. For experiments with B16F10 with the base sgRNA that achieved deep CD47 knockdown, 7 mice received mouse IgG2a isotype control and 13 mice received anti-Tyrp1. All data were collected from three independent experiments. **(B)** Tumor growth curve of projected tumor area versus days after tumor challenge, with B16F10 cells with titrated levels of CD47 and broad expression heterogeneity. Each line represents a separate tumor and is fit with an exponential growth equation: A = A_0_e^kt^. Complete anti-tumor responses in which a tumor never grew are depicted with the same symbol as their growing counterparts and with solid lines at A = 0. A total of 10 mice were challenged with B16F10 with sgRNA^C6G^ (bottom) across two independent experiments, and 5 other mice were challenged with a mosaic B16F10 culture (top) that showed further intermediate levels than the original titrated cell lines. All these mice received anti-Tyrp1 since our *in vivo* studies with B16F10 with deep CD47 knockdown showed no survivors when treated with mouse IgG2a isotype control. All data were collected from 2-3 independent experiments. Inset bar graphs depict response type for each indicated tumor challenge. A partial response was defined as a mouse that survived beyond the median survival of the CD47 deep KD cohort without anti-Tyrp1 (20+ days). **(C)** Survival curves of mice up to 100 days after the tumor challenges in (A) and (B). All mice were challenged with 2×10^5^ B16F10 cells with the listed sgRNA. All tumor inoculations were subcutaneous. For anti-Tyrp1 and mouse IgG2a isotype control treatments, mice were treated either intravenously or intraperitoneally with 250 μg with antibody on days 4, 5, 7, 9, 11, 13, and 15 after tumor challenge. Statistical significance was determined by the Log-rank (Mantel-Cox) test (* p < 0.05). **(D)** Schematic illustrating protocol for sera collection from surviving mice from (A-C) and follow-up experiments to characterize potential *de novo* anti-cancer IgG antibodies. **(E)** (Left) Representative flow cytometry histograms showing that IgG2a/c and IgG2b from convalescent sera binds to both wild-type and Tyrp1 knockout B16F10 cells. (Right) Median fluorescent intensity quantification of IgG2a/c and IgG2b from sera from all surviving mice. Circles represent convalescent sera collected from surviving mice challenged with B16F10 with deep CD47 knockdown (base sgRNA); squares represent sera collected from mice challenged with B16F10 with sgRNA^C6G^. Convalescent sera show statistically significant increase in IgG2a/c titer and a trend in slight increases in IgG2b binding. Binding to even Tyrp1 knockout cells suggests broader recognition of antigens to B16F10. All sera shown were collected between 80-105 days post-challenge. Statistical significance was calculated by an unpaired two-tailed t-test with Welch’s correction (* p < 0.05). **(F)** Phagocytosis of serum-opsonized wild-type, CD47 deep knockdown or CD47/Tyrp1 double knockout B16F10 cells by bone marrow-derived macrophages on two-dimensional tissue culture plastic. Additionally, B16F10 cells treated with either anti-Tyrp1 or mouse IgG2a for opsonization were included as controls for comparisons. Serum IgG derived from survivors retains opsonization and functional phagocytic ability against B16F10. Furthermore, convalescent sera IgG from survivors is still able to drive engulfment of Tyrp1 knockout cells, further suggesting targeting of antigens beyond Tyrp1. Statistical significance was calculated by two-way ANOVA and Tukey’s multiple comparison test between selected groups (mean *±* SD, n = 7 distinct serum samples collected from survivors and n = 6 for non-convalescent serum controls). Circles represent datapoints from sera from survivors challenged with B16F10 with base sgRNA; squares represent survivors from challenges with B16F10 with sgRNA^C6G^.

To further assess how CD47 levels affect higher densities of infiltrating macrophage, we established tumors in mice with B16F10 with wild-type CD47 levels (non-targeting sgRNA), deep repression (base sgRNA), and intermediate CD47-expression levels (sgRNA^C6G^). These mice were then given one intravenous dose of either anti-Tyrp1 or IgG2a control after 4 days. Mice were sacrificed 24 h later, and tumors were harvested, disaggregated, and stained for flow cytometry measurements of immune cell infiltration (**Fig. S7A**). We found that the percentage of CD45+ cells that in the tumor increased with deeper repression when opsonized with anti-Tyrp1 (**Fig. S7B**). Although the overall macrophage percentage of the entire myeloid compartment was similar for all tumors when treated with anti-Tyrp1 (**Fig. S7C**), the overall number of tumor-infiltrating macrophages increased with deeper CD47 repression (**Fig. S7D**). This suggests that macrophages are key effector cells in rejecting initial tumor engraftment when CD47 is depleted.

We then proceeded to collect convalescent serum from survivors to quantify *de novo* anti-cancer IgG generation via acquired immunity. Sera was collected 80 days after the initial challenge and used in antibody binding and 2D phagocytosis assays (**Fig. 3D**). We first quantified pro-phagocytic IgG titer in convalescent sera, focusing on IgG2a and IgG2b, which have been found to engage mouse macrophage Fcγ receptors (Bruhns, 2012; Nimmerjahn et al., 2010). B16F10 cells, either Tyrp1-expresing or Tyrp1 knockout, were incubated with sera (convalescent from survivors or naïve from unchallenged mice) and then counterstained with conjugated antibodies against IgG2a/c and IgG2b (**Fig. 3E**). We found that surviving mice showed significant increases in IgG2a binding in both Tyrp1-positive and Tyrp1 KO B16F10 cells. A similar trend was seen in IgG2b binding, although to a lesser extent. Interestingly though, we find that survivors from deep CD47 knockdown tumors have much higher IgG2a titer than their ∼80% repression (sgRNA^C6G^) counterparts. To test where these *de novo* serum antibodies functionally promote phagocytosis, we performed conventional 2D phagocytosis assays in which cancer cell suspensions were opsonized with sera (or anti-Tyrp1 or mouse IgG2a as controls) (**Fig. 3F**). Cells expressing both CD47 and Tyrp1 were poorly engulfed despite sera nor anti-Tyrp1 opsonization, consistent with the overall inhibitory effect of CD47 on macrophage-mediated phagocytosis. However, under conditions of maximal phagocytosis (CD47-repression and IgG opsonization by anti-Tyrp1), we see that unpurified sera from survivors functionally promotes phagocytosis equivalently to anti-Tyrp1 (∼4-5-fold higher than baseline). We further find that sera continue to promote phagocytosis even in the absence of Tyrp1 in B16F10 CD47/Tyrp1 double knockout cells (∼3-fold higher than baseline), suggesting potent acquired immunity with *de novo* antibodies that target B16F10 antigens beyond Tyrp1.

### Mismatch-CRISPRi titration captures selection of CD47-positive cells in tumor evolution

Given that CRISPRi does not completely repress CD47 and can generate expression heterogeneity, we sought to exploit this weakness to understand how tumor evolution favors CD47-positive cells. In a broad distribution of tumor cells with varying levels of CD47, macrophages may effectively clear low expressors but allow high expressors to escape (Jaiswal et al., 2009), creating a possible opportunity for new emergent cancer population with a selection advantage. We used both *in vitro* tumoroids (**Fig. 4A-i**) and tumors from non-survivors in tumor-challenge experiments (**Fig. 4A-ii**) to assess how CD47 population expression changes after macrophage interaction. After 4 days of co-culture with BMDMs and treatment with anti-Tyrp1, tumoroids showed statistically significant upward shifts in the overall population’s CD47 expression compared to concurrent B16F10-only tumoroid cultures (**Fig. 4B**), suggesting that macrophages clear lower CD47-expressing cells while higher expressers can continue to proliferate. We further found that tumor-derived populations also have more CD47-positive cells compared to their passage-matched *in vitro* control cultures (**Fig. 4C-D; Fig. S8**). This was true for both tumors with sgRNA^C6G^ (∼80% repression) and tumors with base sgRNA (deep repression). FACS-isolated CD47-high cells from these tumor-derived cultures were also more resistant to phagocytic uptake compared to *in vitro* deep knockdown controls in conventional 2D macrophage phagocytosis assays, confirming functional consequences of a tumor population’s overall CD47 recovery.

**Figure 4.**
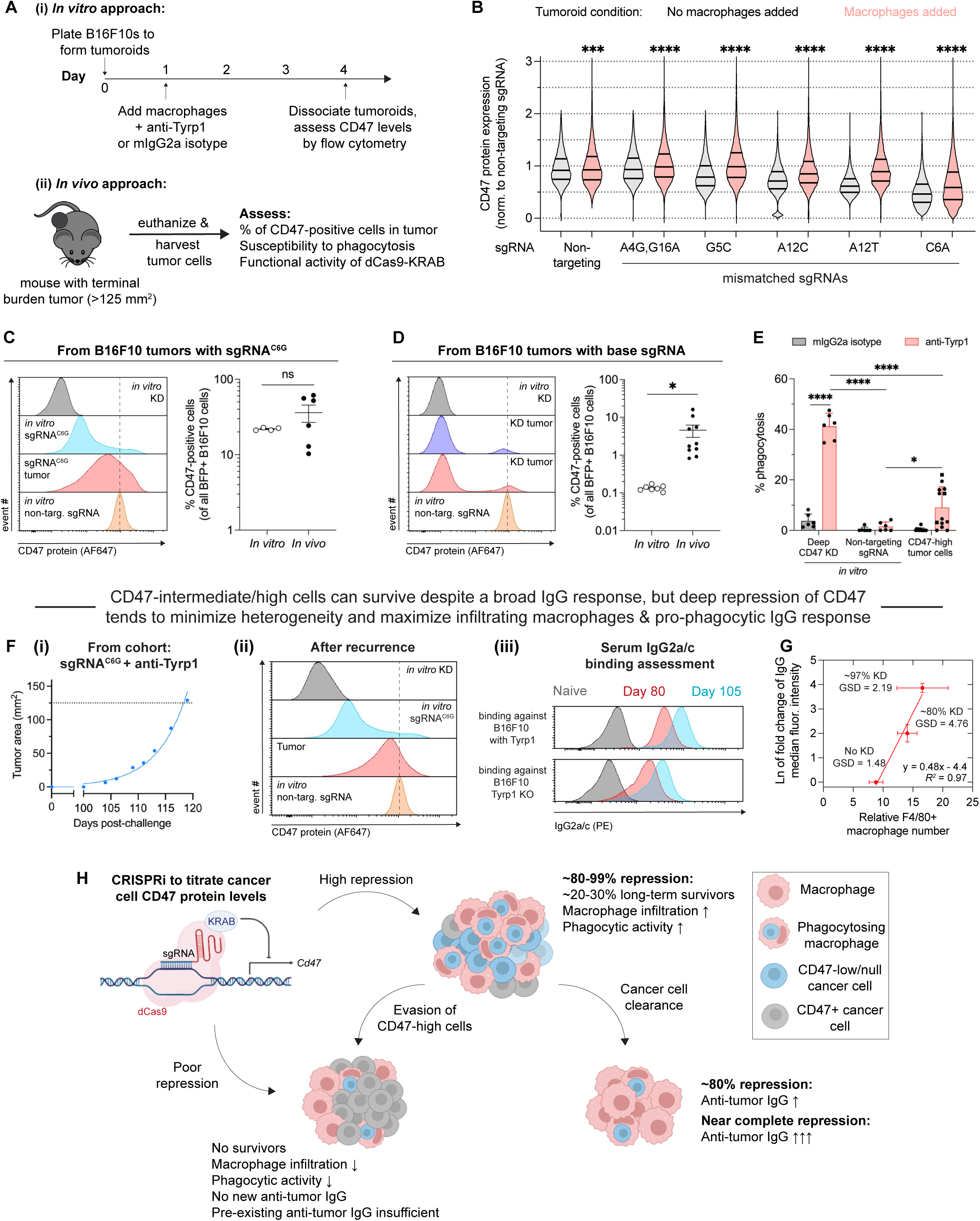
Tumor evolution favors selection and enrichment of CD47-positive cells and can even be responsible for recurrence despite developing acquired immunity. **(A)** (i) Experimental timeline for measuring changes in population CD47 levels in engineered immuno-tumoroids. B16F10 cells were plated on non-adhesive surfaces in a U-bottom shaped well. 24 h later, macrophages were added at a 3:1 ratio to initial B16F10 number, with or without anti-Tyrp1. 72 h after macrophage addition, tumoroids were collected and dissociated for flow cytometry analysis. Concurrently, B16F10 tumoroids with the same sgRNAs but without the addition of macrophages were maintained as a control group. (ii) Experimental timeline for measuring changes in CD47 levels in syngeneic tumor xenografts in C57BL/6 mice. Tumors grew to a terminal burden (∼125 mm^2^), after which mice were humanely euthanized. Tumors were excised and dissociated for flow cytometry analysis and for generating tumor-derived B16F10 cultures for future assays. **(B)** Distribution of CD47 levels in immuno-tumoroids comprised of bone marrow-derived macrophages and B16F10 cells at day 4. All data are normalized to the geometric mean CD47 expression of the concurrent day-matched control tumoroid (no macrophages added) made with B16F10 cells transduced with non-targeting sgRNA. Statistical significance was calculated by the Mann-Whitney test (unpaired, two-tailed, *** p < 0.001, **** p < 0.0001,) comparing the CD47 level distributions between the tumoroid group that received bone-marrow derived macrophages and the group that did not for each sgRNA. **(C-D)** Representative flow cytometry histograms and quantification of CD47 levels in B16F10 cells harvested from terminal burden tumors in Fig. 3A-C. Mean *±* SEM shown, n = 6 tumors generated from B16F10 cells with sgRNA^C6G^ and 4 passage-matched *in vitro* cultures of B16F10 with sgRNA^C6G^; n = 10 tumors generated from B16F10 cells with the base sgRNA and 7 passage-matched *in vitro* cultures of B16F10 with base sgRNA. Statistical significance was calculated by an unpaired two-tailed t-test with Welch’s correction (ns, not significant; * p < 0.05). Dashed line is set at the geometric mean of the non-targeting sgRNA control to facilitate visualization. **(E)** Phagocytosis of B16F10 with non-targeting sgRNA or base sgRNA (for deep CD47 knockdown) and FACS-sorted CD47-high B16F10 cells isolated from tumor-derived cultures in (C-D). Assay was done with bone marrow-derived macrophages, with or without anti-Tyrp1 for opsonization, on two-dimensional tissue culture plastic. For CD47-high tumor cells, circles represent B16F10 cells FACS-sorted from tumors with base sgRNA while squares represent cells from tumors with sgRNA^C6G^. While the tumor-derived CD47-high B16F10 cultures are more readily phagocytosed than *in vitro* B16F10 with non-targeting sgRNA, they are still phagocytosed significantly less than B16F10 cells with deep CD47 knockdown. Statistical significance was calculated by two-way ANOVA and Tukey’s multiple comparison test between selected groups (* p < 0.05; **** p < 0.0001). Mean *±* SD shown, n = 6 distinct cultures for both deep CD47 knockdown and non-targeting sgRNA, n = 13 tumor-derived cultures for CD47-high cells). **(F)** Late cancer recurrence in a mouse from the initial 100-day survivors from the sgRNA^C6G^ cohort. (i) Tumor growth curve for a single mouse that showed cancer recurrence 104 days after the initial challenge and despite the generation of anti-cancer IgG antibodies. The solid line is fit with an exponential growth equation: A = A_0_e^kt^. (ii) Flow cytometry histograms for CD47 protein levels on B16F10 cancer cells harvested from tumor in (F). For comparing CD47 levels, protein levels were also measured on *in vitro* B16F10 cells with the following for comparisons: base sgRNA (deep CD47 knockdown), sgRNA^C6G^, and non-targeting sgRNA. Dashed line is set at the mode of the non-targeting sgRNA control to facilitate visualization. (iii) Flow cytometry histograms and quantification showing that IgG2a/c and IgG2b from the convalescent sera of the mouse that experienced cancer recurrence still binds to both wild-type and Tyrp1 knockout B16F10 cells. Despite active anti-cancer IgG antibody circulating in system, tumor evolution selecting CD47-positive cells can overcome the recently developed acquired immunity. Binding was assessed from convalescent sera collected before cancer recurrence (day 80) and after recurrence (day 105). Dashed line is set at the geometric mean of the non-targeting sgRNA control to facilitate visualization. **(G)** Overall pro-phagocytic IgG binding signal (IgG2a/c + IgG2b, normalized values from naïve sera control) for each sgRNA that produced long-term survivors against the relative tumor-infiltrating macrophage numbers (anti-Tyrp1 treatment conditions from Fig. S6D; all data are originally normalized to macrophage numbers from B16F10 tumors with non-targeting sgRNA treated with mouse IgG2a isotype control). Bottom table contains summary data on knockdown levels and geometric standard deviation for each relevant cell line. **(H)** Summary of experimental findings and proposed CD47-dependent effects on tumor evolution and titer of *de novo* anti-cancer antibodies.

We further confirmed that these tumor-derived cultures still expressed targetable Tyrp1 antigen (**Fig. S9A**) and that any variability in Tyrp1 expression still correlated weakly with the bimodal engulfment behavior observed (**Fig. S9B**). Lastly, we confirmed that KRAB-dCas9 in these tumor-derived cultures was still able to induce deep knockdown (using a Tyrp1-targeting sgRNA for quality control assessment), leading us to conclude that decreased functional activity of the KRAB-dCas9 was unlikely (**Fig. S9C**).

Interestingly, we noted that one mouse from the sgRNA^C6G^ tumor challenge cohort developed a tumor at day 105 (**Fig. 4F-i**). We isolated convalescent sera from this partial responder at this timepoint and again when terminal burden was reached. We harvested the tumor and then assessed the tumor population’s CD47 levels compared to relevant passage-matched *in vitro* controls (**Fig. 4F-ii**). We found that the resulting tumor cells were mostly CD47-high expressors, which illustrates how even a few phagocytosis-resistant cancer cells can ultimately lead to fatal recurrence. Recurrence occurred despite the generation of phagocytosis-promoting anti-cancer IgG antibodies. In fact, IgG2a in particular saw significant increases in titer between days 84 (before recurrence) and day 104 (after recurrence) (**Fig. 4F-ii)**.

After this case of recurrence, we re-evaluated the observation that deep CD47 repression produced the highest pro-phagocytic IgG titer (**Fig. 3E**). Similarly, tumor-infiltrating macrophage numbers also increases with deeper CD47 repression (**Fig. S7D**). An analysis of the correlation between these two parameters suggests that pro-phagocytic IgG titer increases with higher macrophage numbers, which ultimately depends on cancer cell CD47 levels (**Fig. 4G**) that also relates to heterogeneity in levels (**Fig.S2**). This analysis suggests that incomplete disruption of the CD47-SIRPα checkpoint and/or too broad of CD47-expressing heterogeneity with remaining CD47-high cells is sufficient to inhibit overall immune response and dampen acquired immunity.

## Discussion

While CD47 depletion has previously been found to favor tumor rejection and acquired immunity by promoting macrophage-mediated phagocytosis (Dooling et al., 2022; Li et al., 2020), the extent of knockdown/depletion required to achieve therapeutic efficacy is still relatively unknown. In tumoroid models that used low concentrations of anti-CD47 (below 100 nM), we have previously found that even a low remaining amount of available CD47 is sufficient to prevent macrophage-mediated clearance (Dooling et al., 2022). This suggests that there must be a certain threshold of CD47 available for binding to macrophage to SIRPα required to inhibit phagocytosis.

Mismatch-CRISPRi provided an opportunity to generate knockdowns of various degrees while also creating a broad distribution of CD47 expression in cells, allowing us to study CD47 mean expression and variability as variables in determining therapeutic outcome. Therapeutically, this tumor cell CD47 heterogeneity matters in understanding long-term therapeutic outcomes. Even if a bulk population of cells have CD47 levels below this threshold to ultimately stimulate pro-phagocytic activity, CD47-high expressing cells could still escape anti-cancer macrophage activity (**Fig. 4B-D**).

In established B16F10 syngeneic tumors, we find that IgG opsonization drives tumor rejection in cancer cell populations with ∼20% remaining CD47 on average but with overall individual cell remaining CD47 ranging from 4 to 76%. Mice surviving these tumor challenges also produce *de novo* pro-phagocytic IgG antibodies that opsonize B16F10 cells. However, the titer of these *de novo* antibodies in these surviving mice is lower than that in mice who survived B16F10 challenges with near complete CD47 depletion. While only tested thus far in melanoma, these results suggest relevant long-term clinical implications of remaining CD47: Macrophages may have reduced antigen presentation functions because of this small but remaining CD47 surface density, reducing potential phagocytic feedback that leads to anti-tumor IgG. Reduction of anti-tumor IgG then limits acquired immunity mechanisms, allowing a very small percentage of CD47-high or CD47-intermediate to proliferate in the absence of ongoing IgG opsonization treatments, leading to cancer recurrence (**Fig. 4F, 4H**). Altogether, anti-CD47 blockade and/or future *CD47* knockdown delivery approaches are plausible methods for improving therapeutic outcomes, further insight is needed into what “threshold” of CD47 repression or inactivation is needed for favorable outcomes. This study highlights how variability and heterogeneity in CD47 affect therapeutic outcomes and for development of potent acquired immunity.

## Limitations of the study

Our findings pinpoint an interesting critical CD47 regime that determines allows macrophages to dominate tumor proliferation, while also highlighting the importance of single-cell CD47 expression heterogeneity in final therapeutic outcome. Nonetheless, we acknowledge that this threshold may not be the same for all cancer types. In this study, we limited our model to B16F10 melanoma because this cell line expresses Tyrp1 antigen. We and others have found that the available therapeutic monoclonal antibody TA99 can successfully target Tyrp1 and can be used as IgG opsonization to stimulate macrophages (Dooling et al., 2022; Hayes et al., 2020; Khalil et al., 2016). Furthermore, we have found that TA99 can generate cures in a CRISPR/Cas9-mediated *Cd47* knockout B16 model (Dooling et al., 2022), providing us a benchmark from which to begin this study. Other cancer cell lines do not have well characterized monoclonal antibodies that can (1) serve as opsonization or (2) generate cures *in vivo*, which would overall hinder efforts into deciphering CD47 density’s role in modulating phagocytosis. Lastly, we wanted to use a well-controlled and widely used tumor model that is expected to be unaffected by CD47 disruption, since this is representative of CD47 blockade monotherapy showing no clinical efficacy against solid tumors (Horrigan 2017; Jalil et al., 2020). Although B16F10 is a tumor transplant model with the associated limitations, it does not respond to CD47 disruption, unlike other syngeneic models, such as MC38 colon cancer, that respond to CD47 disruption independent of macrophages (Liu et al., 2015; Xu et al., 2017). B16F10 melanoma is also poorly immunogenic in immunocompetent mice, unlike MC38 and CT26 models (Lechner et al., 2013), and although a fraction of human melanomas is durably treated with T cell checkpoint blockade, many human melanomas remain resistant and/or recur.

## Materials and Methods

### Cell culture

B16F10 cells (CRL-6475) were obtained from American Type Culture Collection (ATCC) and cultured at 37°C and 5% CO_2_ in either RPMI-1640 (Gibco 11835-030) or Dulbecco’s Modified Eagle Medium (DMEM, Gibco 10569-010) supplemented with 10% fetal bovine serum (FBS, Sigma F2442), 100 U/ mL penicillin and 100 μg/mL streptomycin (1% P/S, Gibco 15140-122). B16F10 cells were maintained in passage in RPMI-1640 but switched to DMEM at least three days prior to *in vivo* subcutaneous xenografts. 293T human embryonic kidney (CRL-1573) cells were also obtained from ATCC and cultured at 37°C and 5% CO_2_ in DMEM culture media supplemented with 10% FBS and 1% P/S. All cell lines were passaged every 2-3 days when a confluency of ∼80% was reached. For trypsinization, cells were washed once with Dulbecco’s phosphate-buffered saline (PBS, Gibco 14190-136) and then detached with 0.05% Trypsin (Gibco 25300-054) for 5 min at 37°C and 5% CO_2_. Trypsin was quenched with an equal volume of complete culture media.

### Lentiviral production and transduction

24 h prior to lentivirus production, 293T cells were plated at a density of 8×10^5^ cells in 2 mL of DMEM in individual wells of a 6-well plate. On the day of transfection, 1.35 μg of psPAX2, 165 ng of pCMV-VSV-G, and 1.5 μg of lentiviral plasmid were added into a microcentrifuge tube with 7.5 μL of Mirus TransIT-Lenti transfection reagent (Mirus Bio, 6604). Once all plasmids were pooled together with the transfection reagent, 300 μL of serum-free media was added. The solution was gently mixed by pipetting and then allowed to incubate for 30 min. After 30 min elapsed, the solution was gently added dropwise by pipetting to an individual 6-well containing the 293T cells plated the day prior.

For CRISPRi-mediated knockdown of CD47, we used pHR-SFFV-KRAB-dCas9-P2A-mCherry for introducing dCas9 into B16F10 cells and pU6-sgRNA EF1Alpha-puro-T2A-BFP for delivering sgRNAs. Both plasmids were gifts from Jonathan Weissman: Addgene plasmids #60954 and #60955, respectively. For KRAB-dCas9 validation and quality control assessment (Fig. S8C), we used pU6-sgRNA EF1Alpha-puro-T2A-GFP to have a different fluorophore marker for successfully transduced cells. pU6-sgRNA EF1Alpha-puro-T2A-GFP was also a gift from Jonathan Weissman: Addgene plasmid #111596. psPAX2 was a gift from Didier Trono (Addgene plasmid #12260). pVSV-G was a gift from Bob Weinberg (Addgene plasmid #8454).

Viral production was allowed to continue for 48 h, after which the supernatant was collected from each well and centrifuged for 5 min at 300 x *g*. The supernatant was collected and then either added directly to B16F10 cells (0.5 mL of lentivirus-containing supernatant to 2×10^4^ cells in an individual 6-well) or stored at −70°C for long-term storage. 72 h after transduction, spent media with lentivirus was aspirated, B16F10 cells were washed with PBS, and fresh media was added. For lentivirus delivering KRAB-dCas9-P2A-mCherry, successfully transduced B16F10 cells were isolated and enriched to purity via fluorescence-activated cell sorting (FACS) using a BD Biosciences FACSAria Fusion ES. The top ∼30% mCherry-positive cells were FACS-sorted and expanded. For lentivirus delivering sgRNA, fresh media contained 1 μg/mL of puromycin (Invitrogen A1113803) was added. Cells were kept in puromycin-containing until a non-transduced control population also treated with puromycin completely died. We note that we also use B16F10 CD47 and Tyrp1 knockout lines in this study, whose preparation has been previously described (Hayes, et al., 2020; Andrechak et al., 2022).

### Cloning of base and mismatched sgRNAs

All sgRNA oligonucleotides (hereafter simply called oligos) were ordered from Integrated DNA Technology (IDT) and resuspended at 100 μM in Nuclease-Free Duplex Buffer (IDT 11-05-01-03). Top and bottom oligos were annealed in a 50 μL reaction volume containing: 1 μL of each oligo, 23 μL of nuclease-free water, and 25 μL of annealing buffer (200 mM potassium acetate, 60 mM HEPES-KOH (pH 7.4), and 4 mM magnesium acetate). The reaction was incubated at 95°C for 5 min in a PCR machine, and the oligos were then allowed to gradually anneal while cooling to room temperature. The annealed oligos were diluted 1:20 in nuclease-free water and used immediately.

Either pU6-sgRNA EF1Alpha-puro-T2A-BFP or pU6-sgRNA EF1Alpha-puro-T2A-GFP, described originally in (Gilbert et al., 2014), was used for delivery of all sgRNAs in this study. This construct was digested in a 100 μL containing: 6 μg of construct, 10 μL of 10X FastDigest Green Buffer (ThermoFisher B72), 2.5 μL of FastDigest BstXI (ThermoFisher FD1024), 2.5 μL of FastDigest Bpu1102I (ThermoFisher FD0094), and remaining volume nuclease-free water. Linearized vector was run on a 1.0% agarose gel and gel purified using QIAquick Gel Extraction Kit (28704). For ligation, a 20 μL reaction volume was prepared containing: 100 ng of digested vector backbone, 2 μL of the 1:20 diluted annealed oligo, 2 μL of fresh 10X T4 ligase buffer, and 1 μL of T4 DNA ligase (New England BioLabs M0202S). The reaction was incubated for 2 h, after which ligated vector was transformed into DH5α bacteria (ThermoFisher 12297016) using standard heat shock protocols. Individual bacteria colonies were picked and expanded. Plasmid was isolated using QIAprep Spin Miniprep Kit (Qiagen 27104). All plasmids were sent to the Penn Genomic Analysis Core for sgRNA insertion validation by Sanger sequencing. All sgRNAs used are listed in Tables 1 and 2.

**Table 1.**
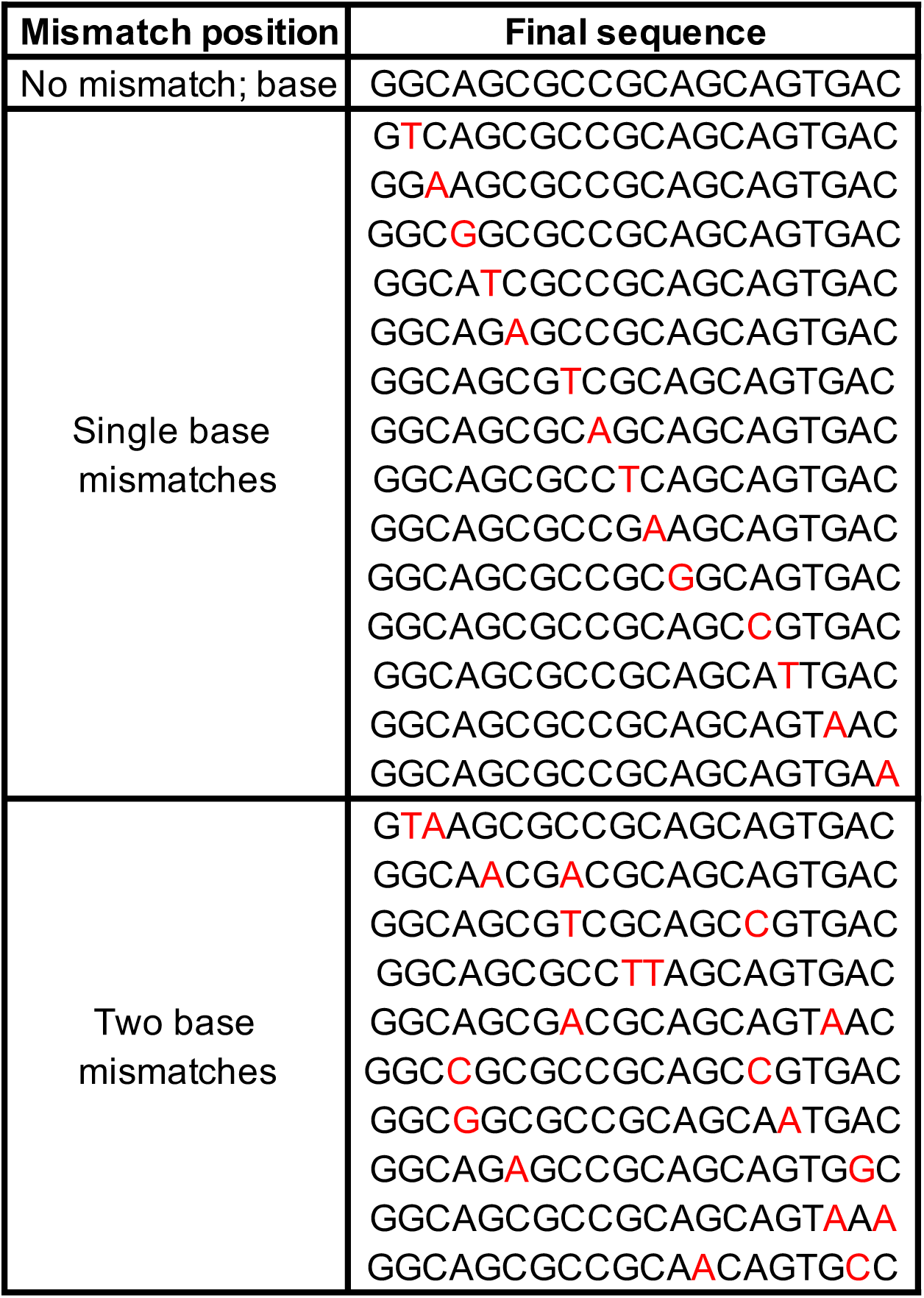
Compact library of mismatches to identify sensitive nucleotide sites.

**Table 2.**
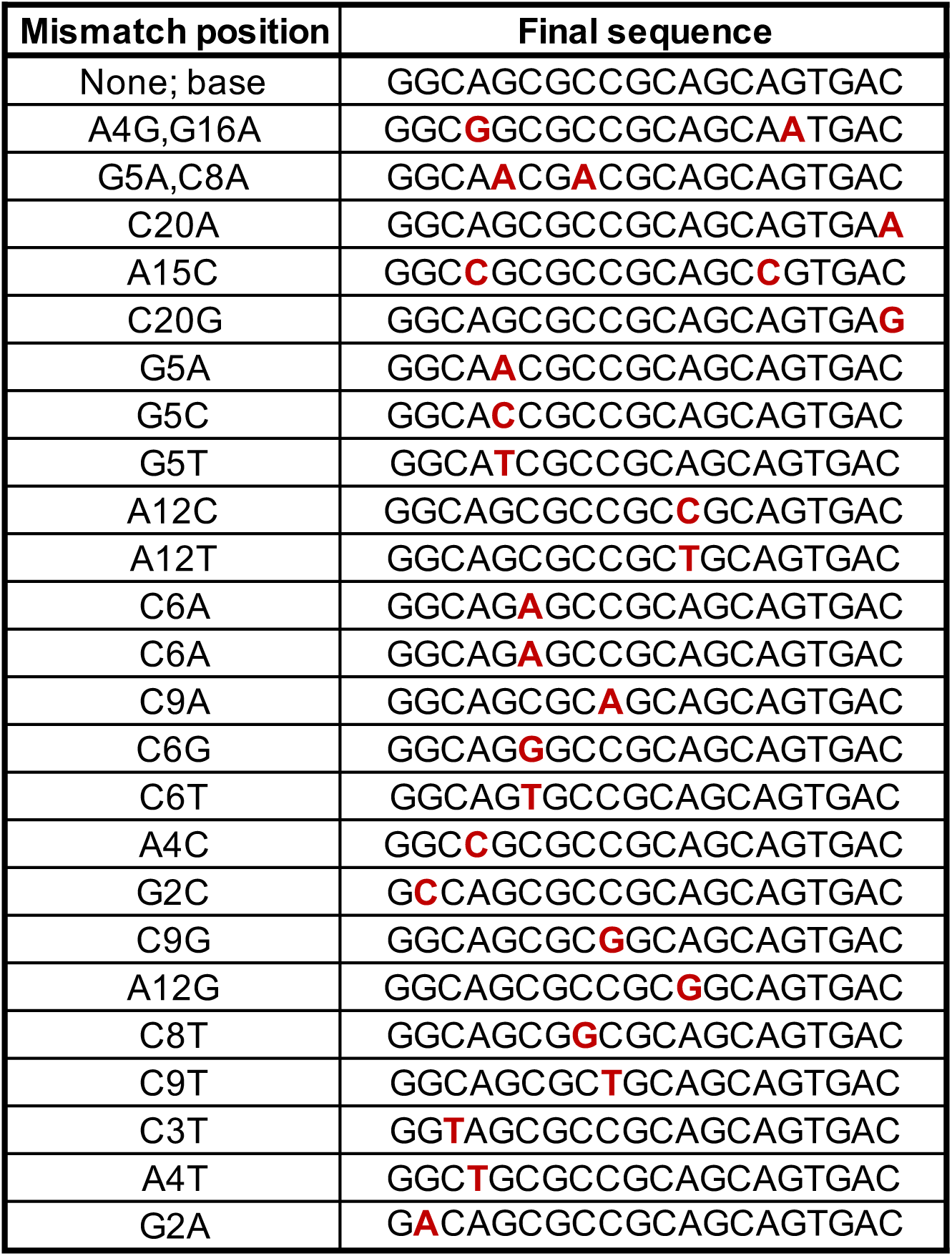
List of final sgRNAs chosen for subsequent experiments.

### Individual evaluation of sgRNAs for knockdown

After B16F10 cells were successfully transduced and puromycin-selected, cultures were briefly trypsinized and then resuspended in 5% (w/v) bovine serum albumin (BSA) dissolved in PBS. Cells were incubated in BSA for 30 min, after which anti-mouse CD47 antibody clone MIAP301 was added to a final concentration of 20 μg/mL. Cells were incubated in primary antibody for 30-45 min on ice. After the incubation period elapsed, cells were centrifuged at 300 x *g* for 5 min, washed once with PBS, centrifuged again at 300 x *g* for 5 min, and then resuspended in 0.1% (w/v) BSA with Alexa Fluor 647 goat anti-rat IgG (see **Antibodies** section for more information). Secondary antibody incubation occurred for 30-45 min, after which cells were centrifuged at 300 x *g* for 5 min, washed once with PBS, centrifuged again at 300 x *g* for 5 min, and then resuspended in 5% (v/v) FBS/PBS. Cells were run on a BD LSRII (Benton Dickinson) flow cytometer. For validation of KRAB-dCas9 in tumor-derived cultures (Fig. S8C), an identical protocol was followed, using anti-Tyrp1 clone TA99 at 20 μg/mL for primary antibody staining of Tyrp1 antigen. The secondary antibody used was Alexa Fluor 647 donkey anti-mouse IgG. Data were analyzed with FCS Express 7 software (De Novo Software). Generation of knockdown distribution histograms and statistics assessment were done using RStudio.

### Antibodies

Antibodies used for *in vivo* treatment and blocking and for *in vitro* phagocytosis are as follows: anti-mouse/human Tyrp1 clone TA99 (BioXCell BE0151), mouse IgG2a isotype control clone C1.18.4 (BioXCell BE0085) Low-endotoxin and preservative-free antibody preparations were used for *in vivo* treatments and *in vitro* phagocytosis experiments. For primary antibody staining of surface proteins via flow cytometry, the following were used: anti-mouse CD47 clone MIAP301 (BioXCell BE0270), anti-mouse/human Tyrp1 clone TA99, anti-mouse SIRPα clone P84 (BioLegend 144004). Secondary antibodies used for flow cytometry are as follows: Alexa Fluor 647 donkey anti-mouse IgG (ThermoFisher A-31571), Alexa Fluor 647 goat anti-rat IgG (ThermoFisher A-21247), Alexa Fluor 488 goat anti-rat igG (ThermoFisher, A-11006). All secondary antibody concentrations used followed the manufacturer’s recommendations.

For immune infiltrate analysis, the following BioLegend antibodies were used: Brilliant Violet 785 anti-mouse CD45 clone 30-F11 (103149), PE/Cyanine7 anti-mouse/human CD11b clone M1/70 (101215), APC anti-mouse F4/80 clone BM8 (123115), PerCP anti-mouse Ly-6G clone 1A8 (127653), Brilliant Violet 605 anti-mouse Ly-6C clone HK1.4 (128035). For IgG titer in tumor challenge surviving mice, the following BioLegend antibodies were used: PE anti-mouse IgG2a clone RMG2a-62 (407108, known to bind IgG2c as well) and APC anti-mouse IgG2b (406711).

### Mice

C57BL/6J mice (Jackson Laboratory 000664) were 6-12 weeks old at the time of tumor challenges and for bone marrow harvesting. All experiments were performed in accordance with protocols approved by the Institutional Animal Care and Use Committee (IACUC) of the University of Pennsylvania.

### Bone marrow-derived macrophages (BMDMs)

Bone marrow was harvested from the femurs and tibia of donor mice, lysed with ACK buffer (Gibco A1049201) to deplete red blood cells, and then cultured on Petri culture dishes for 7 days in Iscove’s Modified Dulbecco’s Medium (IMDM, Gibco 12440-053) supplemented with 10% FBS, 1% P/S, and 20 ng/mL recombinant mouse macrophage colony-stimulating factor (M-CSF, BioLegend 576406). 72 h after initial plating, one whole volume of fresh IMDM supplemented 10% FBS, 1% P/S, and 20 ng/mL M-CSF. After 7 days of differentiation, spent media was removed, BMDMs were gently washed once with phosphate-buffered saline (PBS), and fresh IMDM supplemented 10% FBS, 1% P/S, and 20 ng/mL M-CSF was added.

### *In vitro* phagocytosis

For two-dimensional phagocytosis assays, BMDMs were detached using 0.05% Trypsin and re-plated in either 6-well or 12-well plates, at a density of 1.8×10^4^ cells per cm^2^ in IMDM supplemented 10% FBS, 1% P/S, and 20 ng/mL M-CSF. After 24 h elapsed, BMDMs were labeled with 0.5 μM CellTracker DeepRed dye (Invitrogen C34565), according to the manufacturer’s protocol. Following staining, BMDMs were washed and incubated in serum-free IMDM supplemented 0.1% (w/v) BSA and 1% P/S. B16F10 cells were labeled with either carboxyfluorescein diacetate succinimidyl ester (Vybrant CFDA-SE Cell Tracer, Invitrogen V12883) or CellTracker CMF_2_HC Dye (Invitrogen C12881), also according to the manufacturer’s protocol. B16F10 cells were detached and opsonized with 10 μg/mL anti-Tyrp1, with 10 μg/mL mouse IgG2a isotype control antibody, or 5% (v/v) mouse serum in 1% BSA. Opsonization was allowed for 30-45 min on ice. Opsonized B16F10 suspensions were then added to BMDMs at a ∼2:1 ratio and incubated at 37°C and 5% CO_2_ for 2 h. Non-adherent cells were removed by gently washing with PBS. For imaging, cells were fixed with 4% formaldehyde for 20 min and later imaged on an Olympus IX inverted microscope with a 40x/0.6 NA objective. The Olympus IX microscope was equipped with a Prime sCMOS camera (Photometrics) and a pE-300 LED illuminator (CoolLED) and was controlled with MicroManager software v2. At least 300 macrophages were imaged per individual well for calculation of phagocytosis.

### 3D tumoroid formation and phagocytosis

Tumoroid formation was done following similarly protocols as those used in (Dooling et al., 2022). Briefly, non-TC-treated 96-well U-bottom plates were treated with 100 μL of anti-adherence rinsing solution (StemCell Technologies 07010) for 1 h. The cells were then washed with 100 μL of complete RPMI 1640 cell culture media. This generated surfaces conducive to generating tumoroids and preventing cells from adhering to the well bottom during experiments. B16F10 were detached by brief trypsinization, resuspended at a concentration of 1×10^4^ cells per mL in complete RPMI 1640 cell culture media (10% FBS, 1% P/S) with 50 μM β-mercaptoethanol (Gibco 21985023). 100 μL of this cell suspension was added to each well such that each tumoroid initially started with approximately 1×10^3^ cells. Aggregation of B16F10 cells was confirmed 24 h later by inspection under microcopy. Upon confirmation of tumoroid formation, BMDMs were labeled with 0.5 μM CellTracker DeepRed dye (Invitrogen C34565), according to the manufacturer’s protocol. BMDMs were then detached by brief trypsinization and gentle scraping and resuspended in complete RPMI 1640 cell culture media at a concentration that would allow for delivery of 3×10^3^ BMDMs to each individual tumoroid culture. The BMDM cell suspension was also supplemented with 120 ng/mL M-CSF and antibodies (either anti-Tyrp1 or mouse IgG2a isotype control) such that delivery of 20 μL of this suspension to each individual tumoroid culture result in final concentrations of 20 ng/mL M-CSF and 20 μg/mL of antibody. Tumoroids were imaged on an Olympus IX inverted microscope with a 4x/0.6 NA objective.

For flow cytometry tumoroid experiments (Fig. 4A-I, 4B), B16F10 tumoroids were prepared in the same manner as described above. In addition to co-cultures with BMDMs, separate B16F10 tumoroids with no BMDMs added were maintained concurrently as controls for baseline tumoroid CD47 expression. After 4 days had elapsed for the experiment, tumoroids were dissociated by brief trypsinization and stained as described in the **Individual evaluation of sgRNAs for knockdown** section.

### Tumor models and *in vivo* xenografts

B16F10 cells cultured in DMEM growth media were detached by brief trypsinization, washed twice with PBS, and resuspended at 2×10^6^ cells per mL. Cell suspensions were kept on ice until injection. All subcutaneous injections were performed on the right flank while mice were anesthetized under isoflurane. Fur on the injection site was wet slightly with a drop of 70% ethanol and brushed aside to better visualize the skin. A 100 μL bolus containing 2×10^5^ cells cancer cells was injected beneath the skin. For treatments, mice received either intravenous or intraperitoneal injections of anti-Tyrp1 clone TA99 or mouse IgG2a isotype control clone C1.18.4 (250 μg antibody in 100 μL PBS) on days 4, 5, 7, 9, 11, 13, and 15 post-tumor challenge. Intravenous injections were done via the lateral tail vein. Tumors were monitored by palpation and measured with digital calipers. The projected area was roughly elliptical and was calculated as A = π/4 x *L* x *W*, where *L* is the length along the longest axis and *W* is the width measured along the perpendicular axis. For our studies, a projected area of 125 mm^2^ was considered terminal burden for survival analyses. Mice were humanely euthanized following IACUC protocols if tumor size reached 2.0 cm on either axis, if tumor reached a projected area greater than 200 mm^2^ or if a tumor was ulcerated.

### Serum collection & IgG titer quantification

Blood was drawn retro-orbitally from mice anesthetized under isoflurane, using heparin- or EDTA-coated microcapillary tubes. Collected blood was allowed to clot for 1 h at room temperature in a microcentrifuge tube. The serum was separated from the clot by centrifugation at 1,500 x *g* and stored at −20°C for use in flow cytometry and phagocytosis assays.

For IgG titer quantification, B16F10 cells were detached by trypsinization and incubated with 5% (v/v) mouse serum in 1% BSA. Opsonization was allowed for 30-45 min on ice. After the incubation period elapsed, cells were centrifuged at 300 x *g* for 5 min, washed once with PBS, centrifuged again at 300 x *g* for 5 min, and then resuspended in 0.1% (w/v) BSA with both PE anti-mouse IgG2a clone RMG2a-62 and APC anti-mouse IgG2b (see **Antibodies** section for more information). Anti-IgG conjugated-antibody incubation occurred for 30-45 min, after which cells were centrifuged at 300 x *g* for 5 min, washed once with PBS, centrifuged again at 300 x *g* for 5 min, and then resuspended in 5% (v/v) FBS/PBS. Cells were run on a BD LSRII (Benton Dickinson) flow cytometer.

### Immune infiltrate analysis of tumors

Four days (96 h) post-tumor challenge, mice received a single dose of intravenously delivered anti-Tyrp1 or mouse IgG2a isotype control. 24 later, mice were humanely sacrificed. Tumors were excised and placed into 5% (v/v) FBS/PBS. Tumors were then disaggregated using Dispase (Corning 354235) supplemented with 4 mg/mL of collagenase type IV (Gibco 17104-019) and DNAse I (Millipore Sigma, 101041159001) for 30-45 min (until noticeable disaggregation) at 37°C, centrifuged for 5 min at 300 x *g*, and resuspended in 1 mL of ACK lysis buffer for 12 min at room temperature. Samples were centrifuged for 5 min at 300 x *g*, washed once with PBS, and then resuspended in 5% (w/v) BSA/PBS for 20 min. After 20 min elapsed, fluorophore-conjugated antibodies to immune markers were added to each cell suspension. The following markers were used for analysis: CD45+, CD11b, F4/80, Ly-6C, Ly-6G. Antibody binding occurred for 30 mins while samples were kept on ice and covered from light. Samples were then centrifuged for 5 min at 300 x *g*, washed once with PBS, and resuspended in FluoroFix Buffer (BioLegend, 422101) for 1 h at room temperature prior to analysis on a BD LSRII (Benton Dickinson) flow cytometer. Data were analyzed with FCS Express 7 software (De Novo Software).

### Tumor-derived cultures and evaluation of constitutive knockdown

Mice with terminal tumor burden (>125 mm^2^) were humanely euthanized. Tumors were excised and placed into 5% (v/v) FBS/PBS. Tumors were then disaggregated using Dispase supplemented with 4 mg/mL of collagenase type IV and DNAse I for 30-45 min (until noticeable disaggregation) at 37°C, centrifuged for 5 min at 300 x *g*, and resuspended in 5 mL of ACK lysis buffer for 12 min at room temperature. Samples were centrifuged for 5 min at 300 x *g*, washed once with PBS, and then resuspended in sterile 5% FBS/PBS. Half of the suspension was centrifuged again for 5 min at 300 x *g* and then resuspended in 5% (w/v) BSA/PBS for flow cytometry staining preparation, described in the next paragraph. The remaining half of the total suspension was plated back in 15-cm tissue culture-treated dishes and cultured at 37°C and 5% CO_2_ in RPMI-1640 supplemented with 10% FBS and 1% P/S. The next tumor-derived cultures were passaged, media was further supplemented with 1 μg/mL of puromycin to kill non-B16F10 cells. When all non-B16F10 cells died, cells were detached by brief trypsinization and then resuspended in 5% (w/v) BSA/PBS for flow cytometry staining preparation, described next.

The cell suspension was incubated in 5% BSA/PBS for 5 min, after which anti-mouse CD47 clone MIAP301 was added at a saturating concentration of 20 μg/mL. Cells were incubated for 30-45 min on ice during anti-mouse CD47 primary antibody staining, after which they centrifuged for 5 min at 300 x *g*, washed once with PBS, and then resuspended in sterile 0.1% BSA/PBS containing Alexa Fluor 647 goat anti-rat IgG secondary antibody at a 1:500 dilution of manufacturer provided stock concentration. Cells were again incubated for 30-45 min on ice during secondary antibody staining, after which they centrifuged for 5 min at 300 x *g*, washed once with PBS, and then resuspended in sterile 0.1% BSA/PBS to be run on a flow cytometer.

For the single mouse that experienced cancer recurrence **(Fig. 4E-F)**, the same protocol was used to stain tumor-harvested cells. Here, however, we also split the suspension into two samples to also stain against Tyrp1 with anti-Tyrp1 clone TA99. The secondary antibody used Alexa Fluor 647 donkey anti-mouse IgG.

### Free energy calculations

All free energy and kinetics numerical values were based on the calculations performed using the dashboard application described in (Eslami-Mossallam et al., 2022). Calculated parameters were done using the dashboard’s initial assumptions.

### Statistical analysis and curve fitting

Statistical analyses and curve fitting were performed in GraphPad Prism 9.4. For normalization and histogram generation of large flow cytometry datasets, RStudio 2022.02.3+492 was used. Details for each analysis are provided in the figure legends. Tumor and tumoroid growth curve data (projected area vs time) were fit to the exponential growth model (A = A_0_e^kt^ for tumors and A = A_1_e^k(t-1)^ for tumoroids) using non-linear least squares regression with prefactors A_0_ or A_1_ and *k*, the exponential growth rate. Principal component analysis of different B16F10 cell line with different sgRNAs was performed with Prism 9.4 and RStudio.

### Code availability

Data generated by flow cytometry were analyzed using RStudio 2022.02.3+492 and its standard package ggplot2 for normalization analyses and violin plot generation. For k-means clustering and principal component analysis, R packages tidyverse, cluster, and factoextra were used. No new code central the conclusions of this study was developed.

## Author contributions

Conceptualization: BHH, HZ. Formal analysis: BHH, HZ. Funding acquisition: BHH, JCA, DED. Investigation: BHH, HZ. Methodology: BHH, HZ, JCA. Resources: DED. Visualization: BHH, HZ. Writing: BHH, HZ, DED.

## Acknowledgements

This work was supported by funding from the following sources: NIH R01 HL124106 (DED), U54 CA193417 (DED), NSF GRFP DGE-1845298 (BHH, JCA). The authors acknowledge the following University of Pennsylvania core facilities: Cell Center Stockroom, the Penn Cytomics and Cell Sorting Resource Laboratory, and the Penn Genomic Analysis Core.

## Data availability statement

All data are available within the article and its supplementary information. Data can be provided upon reasonable request from the corresponding author.

## Supplemental Figure Legends

**Supplemental Figure 1.**
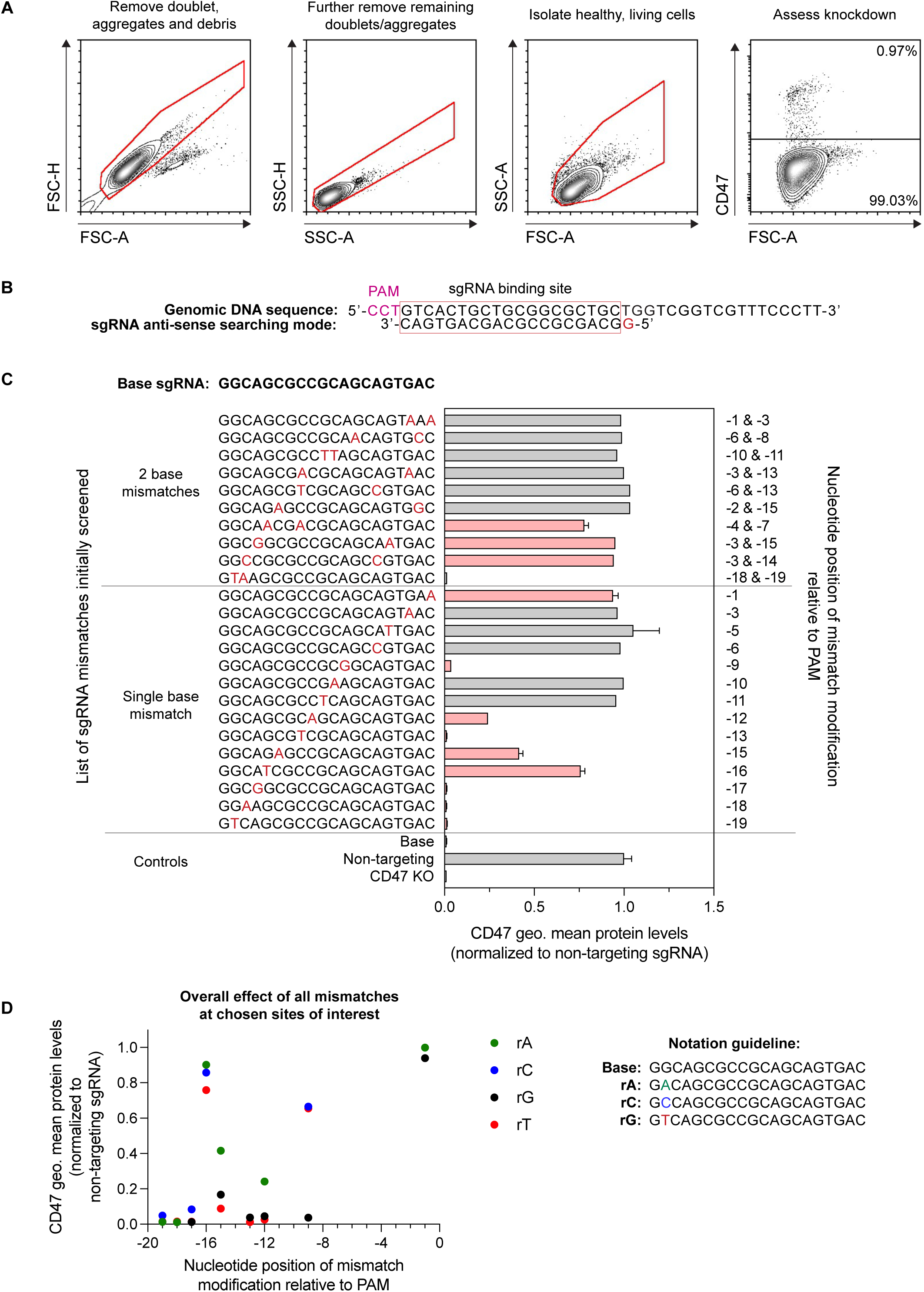
CRISPRi with mismatched sgRNA titrates CD47. **(A)** Representative flow cytometry analysis of lentivirally transduced B16F10 cells to measure CD47 knockdown. Doublet discrimination and dead cell removal was performed with pulse height and pulse area parameters in both the forward (FSC-A vs. FSC-H) and side (SSC-A vs. SSC-H) scatter channels. Healthy, living cells were distinguished from dead cells and debris with the pulse area parameters (FSC-A vs. SSC-A). After lentiviral transduction, B16F10 cells express blue fluorescent protein (BFP). Subsequent gating is done on BFP+ cells only. The flow cytometry contour plots here are an example of CD47 knockdown achieved using the base sgRNA from (Jost et al., 2020). **(B)** Mouse genomic DNA sequence (top) for CD47 that is targeted by the base sgRNA (bottom). The base sgRNA has anti-sense targeting behavior, as illustrated. The protospacer adjacent motif (PAM) is highlighted in magenta. **(C)** Quantification of CD47 knockdown by flow cytometry on a preliminary sgRNA library designed by performing base pair mismatches on the original base sgRNA. Sequences of sgRNAs with their respective mismatches (highlighted in red) are shown on the left. Distance of mismatch relative to the PAM for each sgRNA is shown on the right. We selected working nucleotide positions relative to PAM that produced at least ∼10% knockdown of CD47 levels. Additionally, a B16F10 line (from the same parental line) with CRISPR/Cas9-mediated knockout of CD47 is included for comparison purposes. All data are normalized to the geometric mean CD47 expression of the B16F10 cell line transduced with non-targeting sgRNA. Mean *±* SD shown, n = 2 independent transductions. **(D)** Relative CD47 knockdown levels achieved when performing all possible mismatches at the chosen nucleotide positions relative to PAM, as determined in (C). Additional knockdown levels from the final compact sgRNA library are shown in Fig. 1B.

**Supplemental Figure 2.**
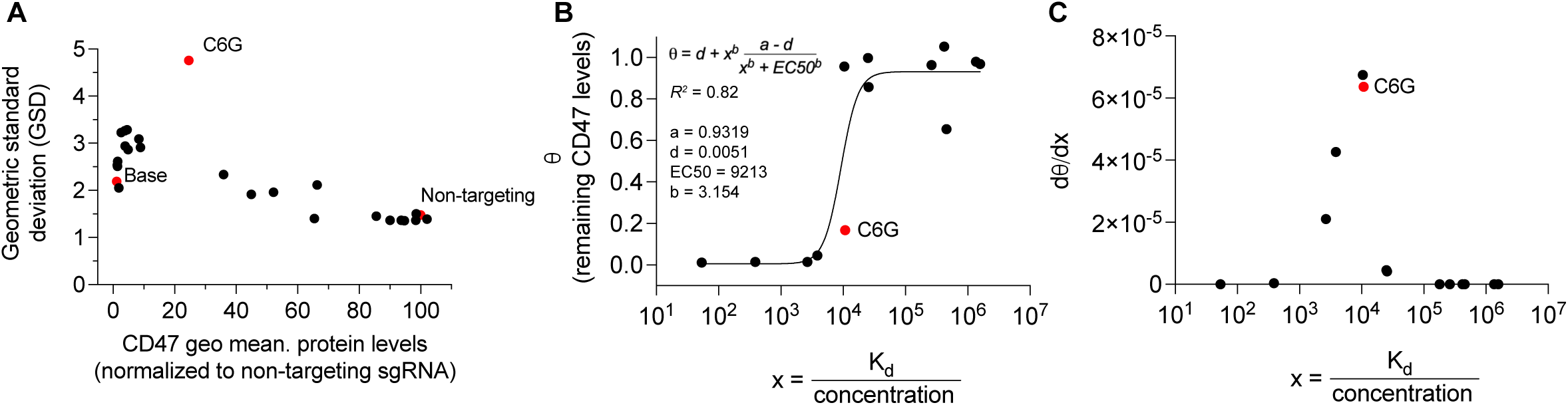
Cell-to-cell variation in expression is highest in the intermediate regime of repression. **(A)** Variation measured by flow cytometry for the range of sgRNAs. **(B)** K_d_ for each sgRNA as calculated from pre-existing modeling software (Eslami-Mossallam et al., 2022) and assuming the same unit ‘concentration’ in each cell for every sgRNA. **(C)** The derivative of (B) shows what is expected for the variation in expression of CD47 for a given sgRNA when the net activity of the sgRNA differs between cells.

**Supplemental Figure 3.**
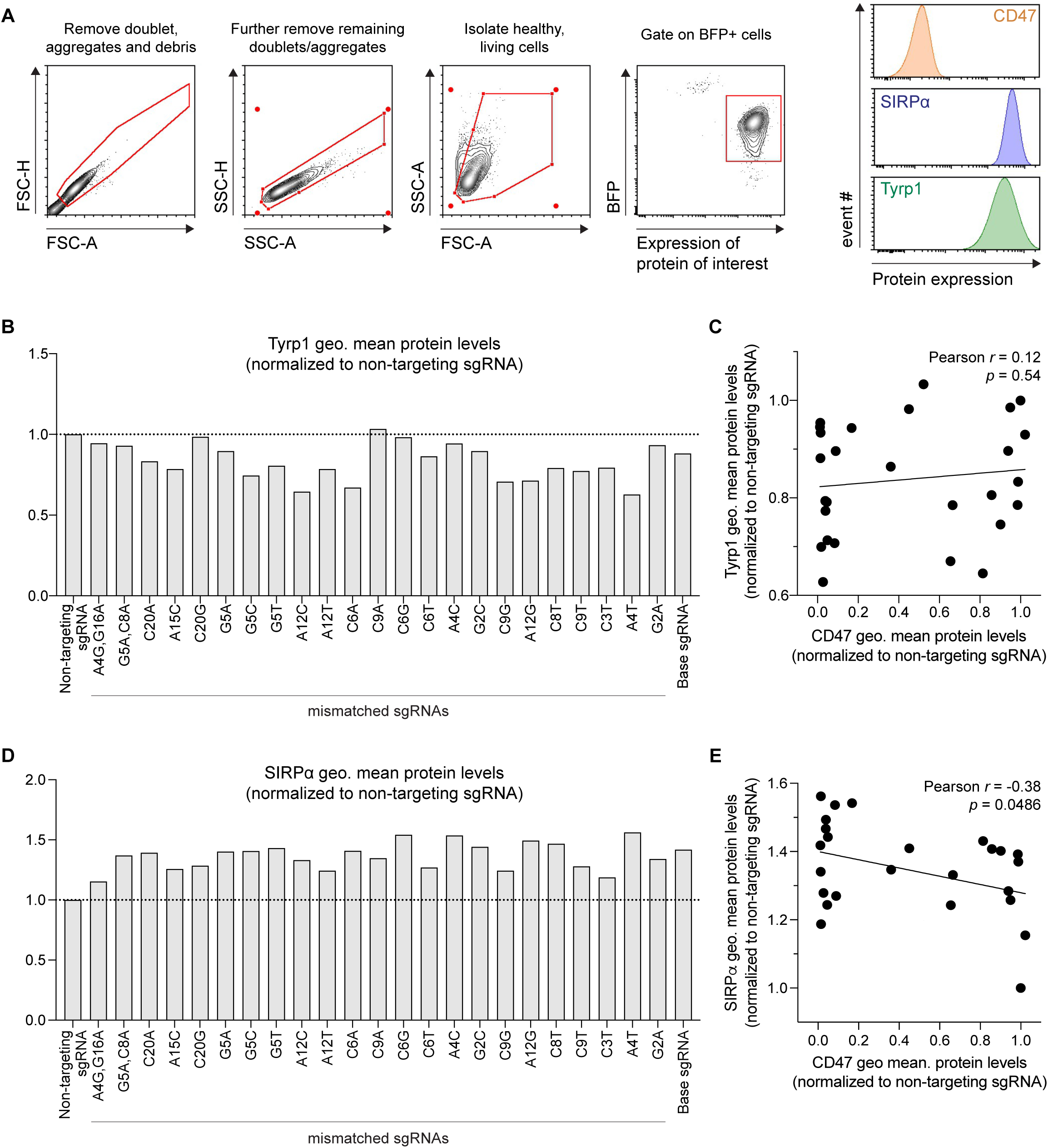
Assessment of Tyrp1 and SIRPα proteins levels after transduction for CD47 titration. **(A)** Representative flow cytometry analysis of lentivirally transduced B16F10 cells to measure both Tyrp1 and SIRPα proteins levels. Doublet discrimination and dead cell removal was performed with pulse height and pulse area parameters in both the forward (FSC-A vs. FSC-H) and side (SSC-A vs. SSC-H) scatter channels. Healthy, living cells were distinguished from dead cells and debris with the pulse area parameters (FSC-A vs. SSC-A). Subsequent gating is then done on BFP+ cells only. The flow cytometry plots here illustrate Tyrp1 and SIRPα proteins levels on B16F10 cells transduced with the base sgRNA (deep CD47 knockdown). **(B)** Quantification of Tyrp1 protein levels in all lentivirally transduced B16F10 cell lines. All data are normalized to the geometric mean Tyrp1 expression of the B16F10 cell line transduced with non-targeting sgRNA. **(C)** Tyrp1 protein levels correlated (Pearson *r* = 0.12, p = 0.54) with the percentage of CD47 knockdown (relative to non-targeting sgRNA). Analysis shows poor correlation between CD47 knockdown and final Tyrp1 levels, suggesting lentiviral transduction has little effect on Tyrp1 expression. **(D)** Quantification of SIRPα protein levels in all lentivirally transduced B16F10 cell lines. All data are normalized to the geometric mean SIRPα expression of the B16F10 cell line transduced with non-targeting sgRNA. **(E)** SIRPα protein levels correlated (Pearson *r* = −0.38, p = 0.0486) with the percentage of CD47 knockdown (relative to non-targeting sgRNA). Analysis shows some correlation between CD47 knockdown and final SIRPα levels, suggesting repression of CD47 reduces *cis* interactions between CD47 and SIRPα on the B16F10 cell surface, freeing availability of SIRPα protein on the cell surface.

**Supplemental Figure 4.**
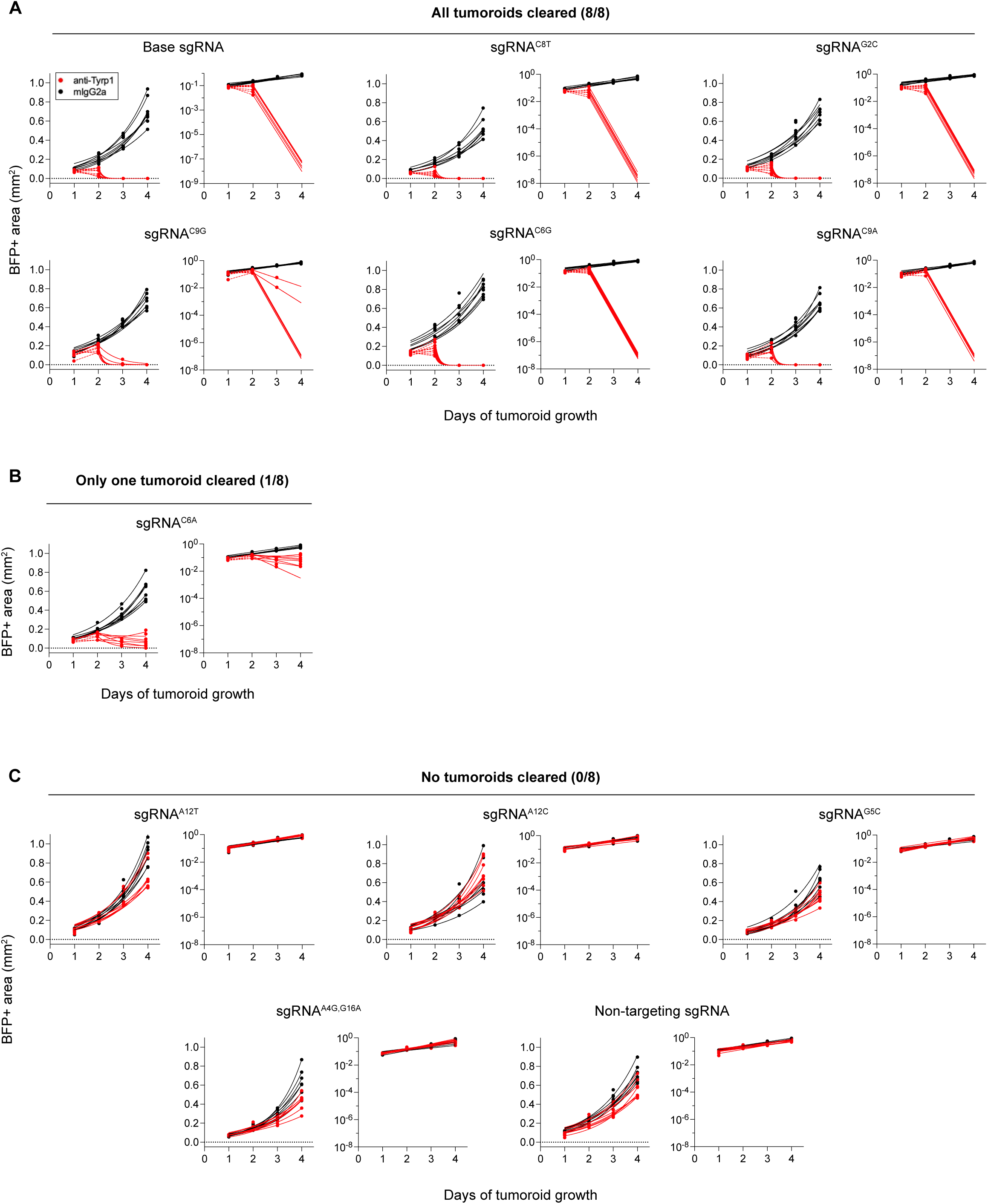
Individual growth curves for all tumoroids. Individual tumoroid growth for all individual tumoroids was measured by calculating the BFP+ area at the indicated timepoints (mean *±* SD, n = 8 total tumoroids from two independent experiments for each sgRNA). Each cell line with a unique sgRNA tested in Fig. 2D is shown, with the left plot of the corresponding sgRNA showing growth on a linear scale and the right plot showing growth on a logarithmic scale. **(A)** Individual growth curves for all B16F10 tumoroids with sgRNAs that resulted in complete clearance. All logarithmic scale plots show downward trends in tumoroid area when treated with anti-Tyrp1, further supporting complete clearance. **(B)** Individual growth curves for B16F10 tumoroids with sgRNA^C6A^, which only resulted in 1/8 of tumoroids being cleared and ultimately displayed plateaued clearance (shown in logarithmic scale plot). **(C)** Individual growth curves for all B16F10 tumoroids with sgRNAs that resulted in no clearance at all. All logarithmic scale plots show upward trends in tumoroid area despite anti-Tyrp1 treatment, suggesting low macrophage phagocytic ability that cannot compete with cancer cell proliferation.

**Supplemental Figure 5.**
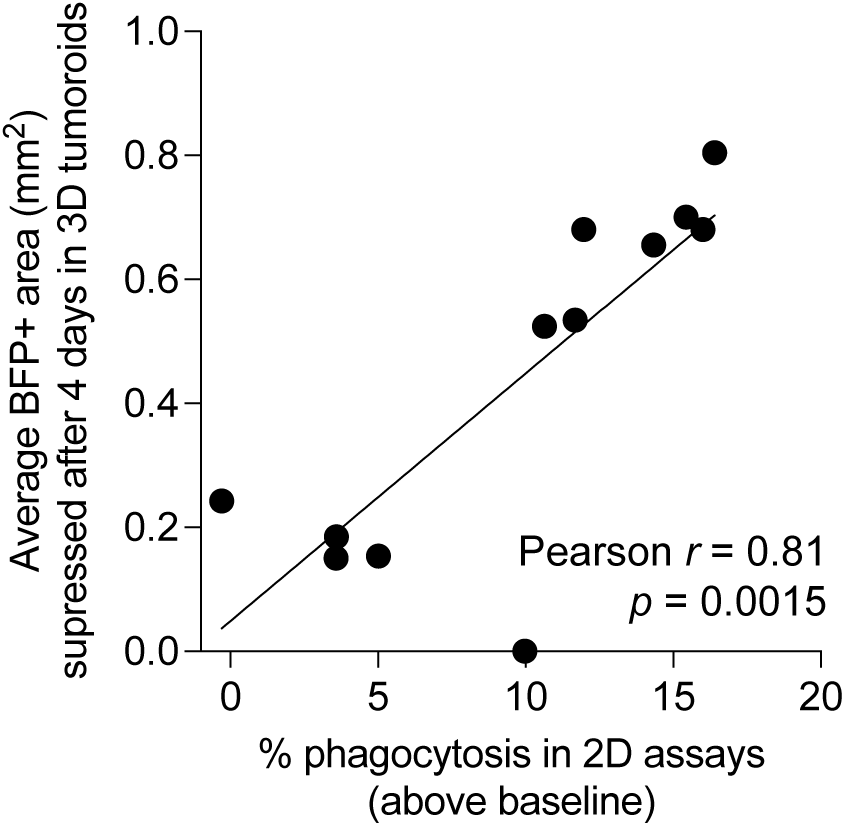
2D phagocytosis assays predict 3D tumoroid clearance for the respective CD47 knockdown. Pearson correlation (Pearson *r* = 0.94, p < 0.001) between 3D tumoroid proliferation potential (area, in mm^2^) and 2D phagocytic activity of macrophages. The percentage of phagocytosis in 2D assays used here subtracts the baseline phagocytic activity of macrophages (mouse IgG2a isotype control for opsonization) from the phagocytic activity when anti-Tyrp1 is used for opsonization to determine the overall contribution of proper IgG opsonization. For tumoroids, the difference in BFP+ area at day 4 between the mouse IgG2a isotype group and the anti-Tyrp1 group was calculated to assess the overall impact macrophages had on inhibiting 3D proliferation.

**Supplemental Figure 6.**
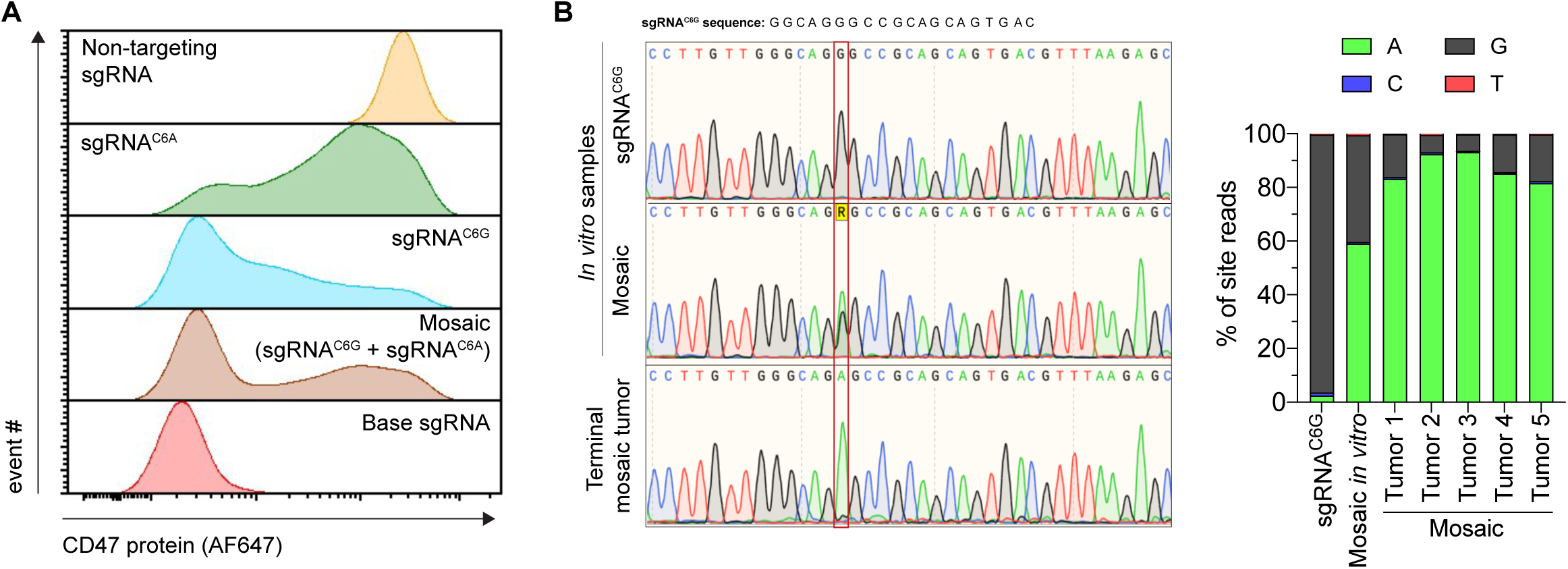
Artificially generated mosaic B16F10 CD47 knockdown cell line highlights *in vivo* selection of CD47-positive cells during tumor evolution. **(A)** Representative flow cytometry histograms for validating the mosaic B16F10 CD47 knockdown cell line. The mosaic cell line was made by mixing B16F10 cells harboring either sgRNA^C6G^ or sgRNA^C6A^ at approximately a 1:1 ratio. **(B)** (Left) Representative Sanger sequencing results for validating the mosaic B16F10 line and for confirming selection of the dominant subpopulation after *in vivo* growth. Red frame highlights nucleotide position of base pair mismatch. (Right) Quantification of Sanger sequencing reads at the nucleotide position of base pair mismatch. The five mosaic B16F10 samples were derived from excised terminal burden tumors subcutaneously xenografted in C57BL/6 mice. All tumor-derived samples show significant increases in the number of adenine (A) reads at the nucleotide position of the mismatch, suggesting that CD47+ cells in the mosaic tumor can more readily proliferate compared to their CD47-low counterparts that are actively engulfed by macrophages.

**Supplemental Figure 7.**
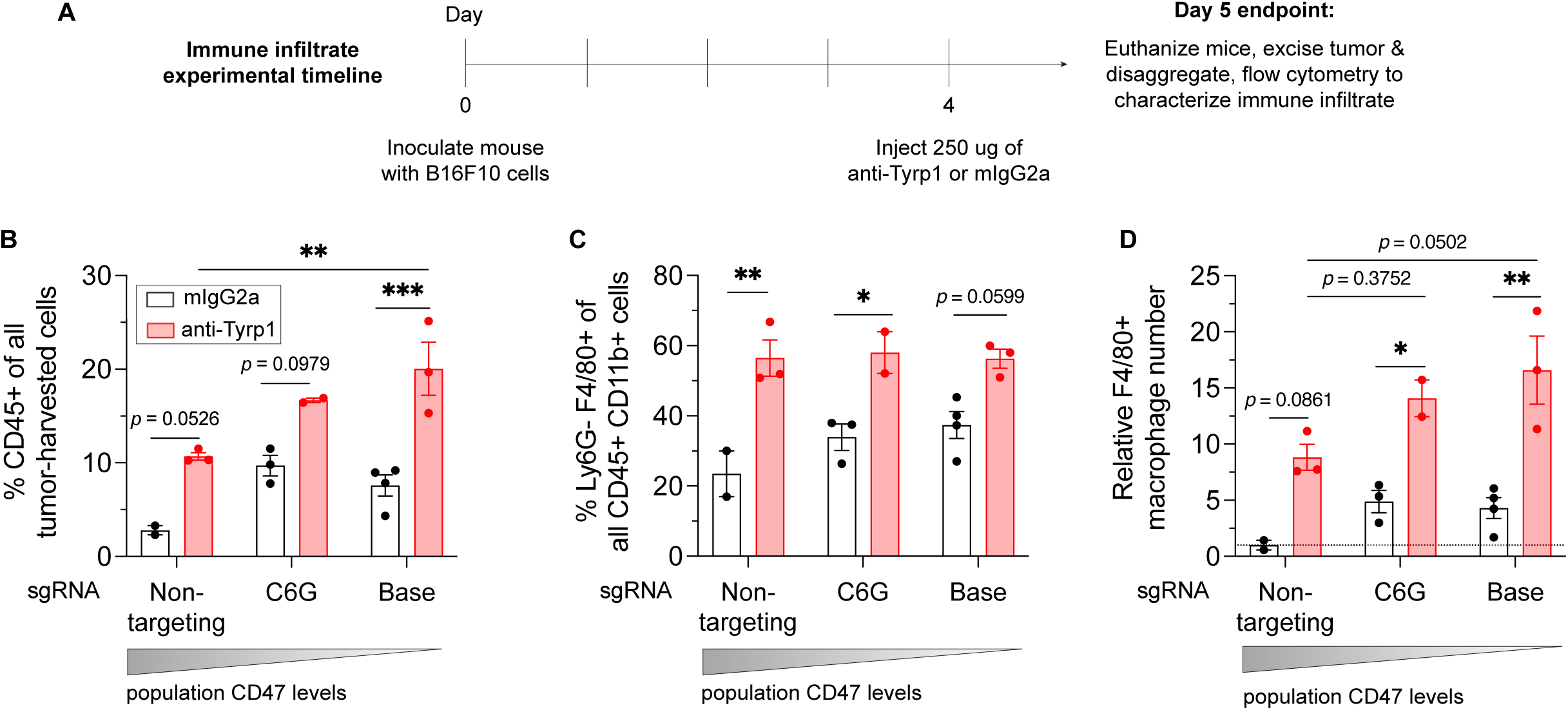
Immune cell infiltration increases in tumors as CD47 levels decrease and with therapeutic opsonization by IgG. **(A)** Experimental timeline for immune cell infiltration analyses in B16F10 tumors. 2×10^5^ B16F10 cells with the listed sgRNA were subcutaneously injected into C57BL/6 mice. Four days (96 h) post-challenge, mice were treated with 250 μg of anti-Tyrp1 or mouse IgG2a isotype. Mice were then euthanized 24 h after antibody treatment, and their tumors were excised and disaggregated for immune infiltrate analysis by flow cytometry. **(B)** Quantification of CD45+ (immune) cells in the excised tumors. In addition to B16F10 with non-targeting sgRNA, Only B16F10 cells harboring sgRNA^C6G^ and base sgRNA were used for immune infiltrate analyses since these were the only lines that had survivors in tumor growth and survival experiments (Fig. 3A-C). Regardless of B16F10 CD47 levels, a single dose of anti-Tyrp1 four days post-challenge showed a trend of a higher number of immune cells in the tumor. However, the overall percentage of immune cells among all tumor cells increased as CD47 was repressed. Statistical significance was calculated by two-way ANOVA and Tukey’s multiple comparison test between selected groups (mean *±* SEM, n = 2-4 tumors per condition, ** p < 0.01, *** p < 0.001). **(C)** Quantification of Ly6G- F4/80+ immune cells among all myeloid cells in the excised tumor. This immune cell marker combination is indicative of macrophages. A single dose of anti-Tyrp1 increased the percentage of macrophages in the entire myeloid population, regardless of the B16F10 CD47 levels at the time of challenge. No statistical difference was noted in macrophage composition of the myeloid cells between tumors of different sgRNAs when treated with anti-Tyrp1. Statistical significance was calculated by two-way ANOVA and Tukey’s multiple comparison test between selected groups (mean *±* SEM, n = 2-4 tumors per condition, * p < 0.05, ** p < 0.01). **(D)** Quantification of relative tumor-infiltrating macrophage numbers from each tumor. Values were calculated by multiplying the percentage of CD45+ cells by the percentage of F4/80+ cells. The product was then normalized to the value from B16F10 tumors with non-targeting sgRNA and treated with mouse IgG2a isotype control (represented by a dash line). Although in (C) we saw no statistically significant change in the macrophage composition of the myeloid compartment, the overall increase in CD45+ cells from (B) indicate that tumors with increased repression of CD47 show a trend of increasing tumor-infiltrating macrophages when treated with IgG opsonization. Statistical significance was calculated by two-way ANOVA and Tukey’s multiple comparison test between selected groups (mean *±* SEM, n = 2-4 tumors per condition, * p < 0.05, ** p < 0.01).

**Supplemental Figure 8.**
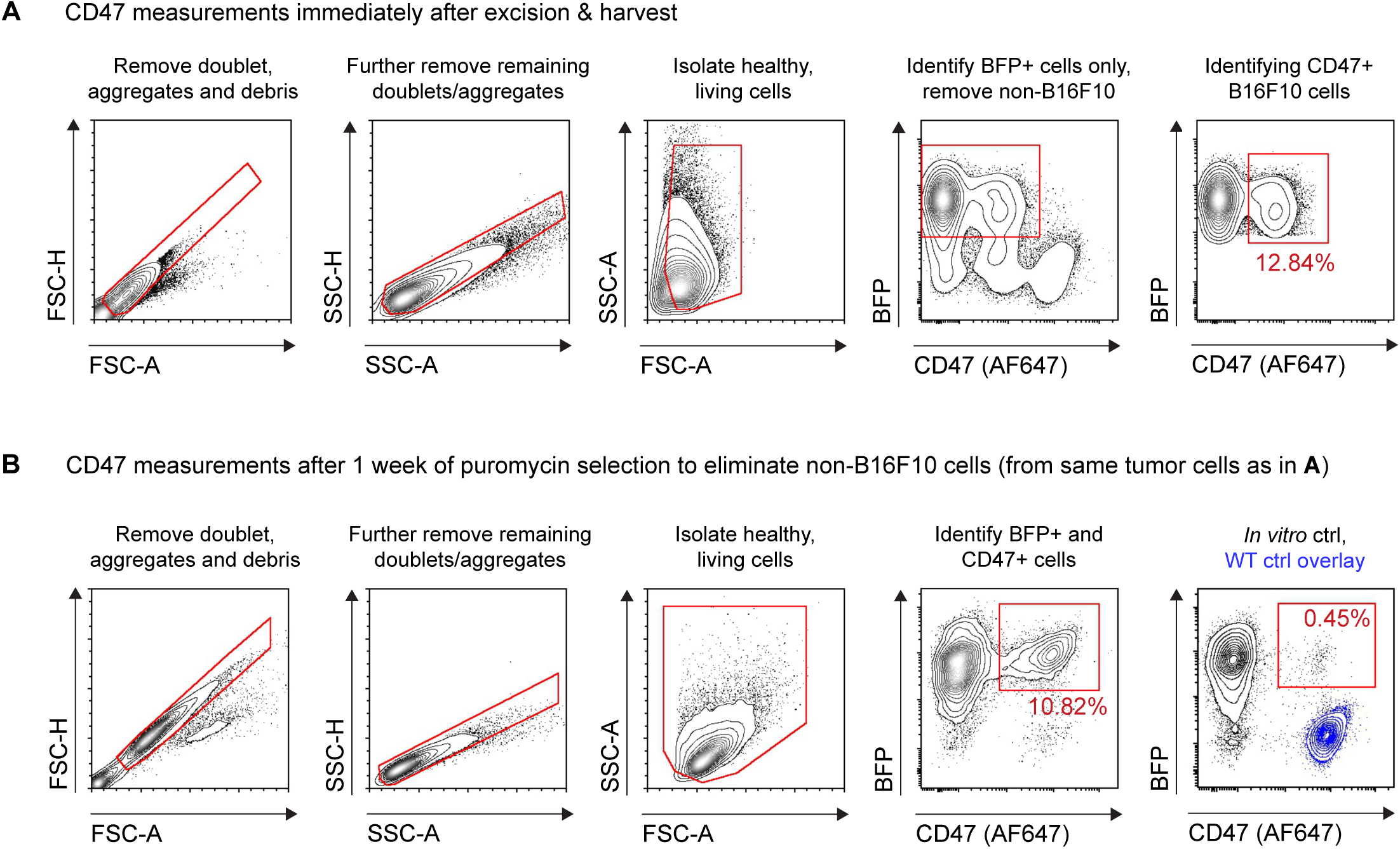
Flow cytometry gating strategies for isolation and analysis of CD47-positive cells from terminal burden tumors. Representative flow cytometry analyses of B16F10 cells harvested from a terminal burden tumor. Doublet discrimination and dead cell removal was performed with pulse height and pulse area parameters in both the forward (FSC-A vs. FSC-H) and side (SSC-A vs. SSC-H) scatter channels. Healthy, living cells were distinguished from dead cells and debris with the pulse area parameters (FSC-A vs. SSC-A). Subsequent gating is then done on BFP+ cells only. **(A)** After gating on healthy cells, we proceeded to gate only on BFP+ cells for a preliminary assessment of CD47 levels on the B16F10 cells to determine if CRISPRi repression was constitutive *in vivo*. We found that many tumors showed a significant subpopulation of CD47-positive cells. **(B)** Tumor-harvested cells were cultured in puromycin for one week to remove non-B16F10 cells. Flow cytometry was repeated, with gating done on BFP+ cells only. Like the results immediately post-harvest, we found that there was still a significant subpopulation of CD47-positive cells in the tumor-derived cultures, even one week after puromycin selection. These results suggest that tumors are rapidly enriched for CD47-positive cells, highlighting relevant vulnerabilities that incomplete CD47 depletion can have on therapeutic outcomes.

**Supplemental Figure 9.**
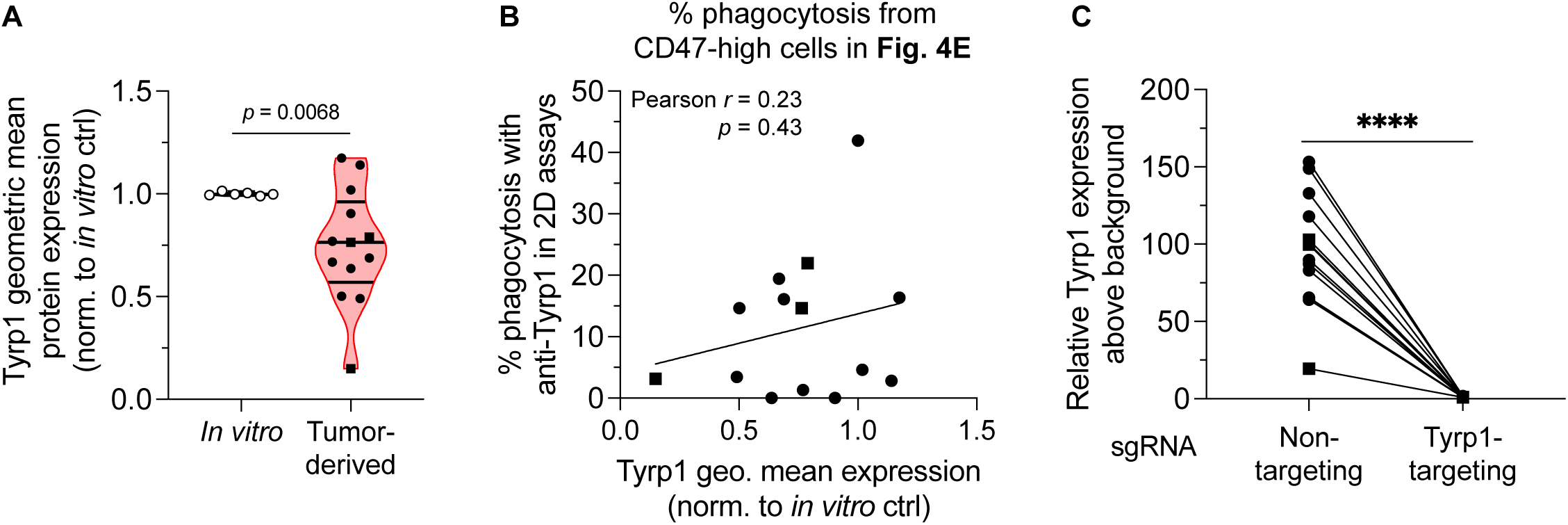
Quality control assessment shows tumor-derived cultures still express Tyrp1 and have functional KRAB-dCas9. **(A)** Quantification of Tyrp1 geometric mean protein expression from tumor-derived cultures from Fig. 4C-D and *in vitro* B16F10 controls. Tumor-derived cultures show a range of Tyrp1 levels but do not show selection of Tyrp1-negative populations. Statistical significance was calculated by an unpaired two-tailed t-test with Welch’s correction (n = 6 for *in vitro* control, n = 13 for tumor-derived cultures; circles represent tumor-derived cultures from B16F10 cells with base sgRNA while squares represent tumor-derived cultures from B16F10 cells with sgRNA^C6G^. **(B)** Pearson correlation analysis (Pearson *r* = 0.23, p < 0.43) between phagocytosis activity in Fig. 4E and Tyrp1 geometric mean levels do not show a statistically significant correlation. **(C)** Quantification of Tyrp1 levels in tumor-derived cultures, lentivirally transduced with non-targeting sgRNA or Tyrp1-targeting sgRNA. Tyrp1 expression is normalized to anti-Tyrp1 binding on B16F10 Tyrp1 knockout cells. After transduction with Tyrp1-targeting sgRNA, tumor-derived cultures show near complete repression of Tyrp1 antigen, suggesting that the KRAB-dCas9 is still functionally active in all cells. This further suggests that the generation of CD47-positive populations in tumors is generated through selection and/or other complex mechanisms. Statistical significance was calculated by an unpaired two-tailed t-test with Welch’s correction (for each condition, n = 13 for tumor-derived cultures; circles represent tumor-derived cultures from B16F10 cells with base sgRNA while squares represent tumor-derived cultures from B16F10 cells with sgRNA^C6G^.

